# Reconstitution of cytolinker-mediated crosstalk between actin and vimentin

**DOI:** 10.1101/2023.10.27.564417

**Authors:** Irene Istúriz Petitjean, Quang D. Tran, Angeliki Goutou, Zima Kabir, Gerhard Wiche, Cécile Leduc, Gijsje H. Koenderink

**Affiliations:** Department of Bionanoscience, Kavli Institute of Nanoscience Delft, Delft University of Technology, 2629 HZ Delft, The Netherlands; Université Paris Cité, CNRS, Institut Jacques Monod, F-75013 Paris, France; Department of Biochemistry and Cell Biology, Max Perutz Laboratories, University of Vienna, Vienna, Austria

**Keywords:** Plakins, Plectin, Cytoskeleton, Rheology, Binding kinetics

## Abstract

Cell shape and motility are determined by the cytoskeleton, an interpenetrating network of actin filaments, microtubules, and intermediate filaments. The biophysical properties of each filament type individually have been studied extensively by cell-free reconstitution. By contrast, the interactions between the three cytoskeletal networks are relatively unexplored. They are coupled via crosslinkers of the plakin family such as plectin. These are challenging proteins for reconstitution because of their giant size and multidomain structure. Here we engineer a recombinant actin-vimentin crosslinker protein called ‘ACTIF’ that provides a minimal model system for plectin, recapitulating its modular design with actin-binding and intermediate filament-binding domains separated by a coiled-coil linker for dimerisation. We show by fluorescence and electron microscopy that ACTIF has a high binding affinity for vimentin and actin and creates mixed actin-vimentin bundles. Rheology measurements show that ACTIF-mediated crosslinking strongly stiffens actin-vimentin composites. Finally, we demonstrate the modularity of this approach by creating an ACTIF variant with the intermediate filament binding domain of Adenomatous Polyposis Coli. Our protein engineering approach provides a new cell-free system for the biophysical characterization of intermediate filament-binding crosslinkers and for understanding the mechanical synergy between actin and vimentin in mesenchymal cells.

**HIGHLIGHTS:**

- Engineering of a recombinant actin-vimentin crosslinker called ACTIF with the plectin intermediate filament binding domain (IFBD), calponin homology domains that mediate actin binding, and a coiled-coil linker.
- ACTIF crosslinks F-actin and vimentin and mediates their co-localization *in vitro*.
- ACTIF has a binding affinity for vimentin that is about 500 times higher than for F-actin.
- ACTIF forms composite bundles of F-actin and vimentin filaments.
- Composite F-actin/vimentin networks stiffen upon crosslinking with ACTIF.
- The design of actin-vimentin crosslinker is modular, as other IFBDs like APCn2 can also be used.

## 1. INTRODUCTION

The cytoskeleton serves multiple crucial functions within mammalian cells. It defines cell shape (Fletcher and Mullins 2010), provides mechanical stability (Pegoraro et al. 2017), drives cell movement and division (Nakaseko and Yanagida 2001), connects cells within tissues (Hoelzle and Svitkina 2012), and influences intracellular signaling processes (Janmey 1998). The cytoskeleton is a network of three primary types of biopolymers: actin filaments (F-actin), microtubules (MTs), and intermediate filaments (IFs), each playing distinct roles in various cellular activities. F-actin and MTs have been widely studied for their roles in regulating cell mechanics and essential processes such as cell division and migration (Dogterom and Koenderink 2019). Both biopolymers form dynamic networks that can generate forces by active (de)polymerization and by the activity of molecular motor proteins. F-actin networks drive different modes of cell migration and also drive membrane constriction in dividing cells by forming a contractile ring. At the same time, F-actin is an important determinant of cell stiffness and strength (Fletcher and Mullins 2010). MTs build the mitotic spindle that drives chromosome segregation in dividing cells, while also contributing to the overall mechanical integrity of the cytoskeleton (Dogterom and Koenderink 2019). By contrast, much less is known about the role of IFs in cell function. Traditionally, they are considered mostly as a kind of “cellular safety belt” that protects cells against large mechanical stresses (Hu et al. 2019, Janmey et al. 1991, Pogoda et al. 2022). Indeed IFs can stretch to more than 3 times their rest length because they are built up of fibrous subunits that can unfold (Block et al. 2017, Forsting et al. 2019). However, in the past years, many additional non-mechanical functions of IFs have been discovered (Patteson et al. 2020). IF networks contribute to the functioning of several cellular organelles (Chernoivanenko et al. 2015, Styers et al. 2004) and regulate signaling pathways that control cell survival, growth, and differentiation (Ridge et al. 2022, Sjöqvist et al. 2021). Intriguingly, the IF protein family comprises 70 different proteins whose expression pattern is cell-type-specific and changes during development and in disease. This raises the interesting possibility that IFs confer cell-type-specific functions. The two most well-studied IFs are keratins, expressed in non-motile epithelial cells, and vimentin, expressed in motile mesenchymal cells. Recent biophysical studies indicate that keratin and vimentin have distinct mechanical properties tailored to their cellular functions (Lorenz et al. 2023).

The three cytoskeletal biopolymers have traditionally been regarded as independent systems with separate cellular tasks. However, growing evidence shows that their functions are coordinated and that this coordination is essential for many core cellular functions (Huber et al. 2015). Directional cell migration, for instance, relies on F-actin to generate the driving force, but coordination with both MTs and IFs is required to polarize the F-actin cytoskeleton and control the amplitude and directionality of force generation (Costigliola et al. 2017, Gan et al. 2016, Jiu et al. 2015, 2017, van Bodegraven and Etienne-Manneville 2020). In dividing cells, F-actin/MT interactions control the correct placement of the division plane and F-actin/IF interactions at the cell cortex are essential for normal mitotic progression (Duarte et al. 2019, Serres et al. 2020). These examples underscore the need to study the mechanisms that couple the activities of the three cytoskeletal systems.

Cytoskeletal coupling is known to involve a combination of biochemical coupling via signaling loops and transcriptional regulation (Tomar and Schlaepfer 2009) and physical coupling via crosslinking proteins (Prechova et al. 2022), motor proteins (Seetharaman and Etienne-Manneville 2020), and joint connections to adhesion receptor complexes in the plasma membrane (Osmanagic-Myers et al. 2015) and to LINC complexes in the nuclear envelope (Tusamda Wakhloo et al. 2020). This complexity makes it difficult to understand the mechanisms of cytoskeletal coupling in cells. Cell-free reconstitution can provide a way to overcome this challenge. In this reductionist approach, components of the cytoskeleton are purified and studied in isolation or together with a limited set of regulatory proteins. In the past, cell-free reconstitution of isolated cytoskeletal biopolymers has successfully delivered quantitative models explaining how F-actin drives cell motility (Blanchoin et al. 2014), how MTs organize the mitotic spindle (Vleugel et al. 2016), and how IFs provide mechanical resilience (Rölleke et al. 2023). However, coupling between different cytoskeletal subsystems has only recently begun to be investigated by reconstitution. Most of the available work has focused on F-actin/MT coordination, with studies of molecular motor motility at F-actin-MT crossings (Ali et al. 2007, Schroeder et al. 2010), interdependent F-actin/MT polymerization (Alkemade et al. 2022, Breitsprecher et al. 2012, López et al. 2014), and mechanical synergy in F-actin/MT composites (Pelletier et al. 2009, Ricketts et al. 2018, Sheung et al. 2021). There are even fewer studies of cytoskeletal composites involving IFs. A few studies have investigated the collective viscoelastic properties of two-component (IF/F-actin) (Esue et al. 2006, Golde et al. 2018, Jensen et al. 2014) and three-component composite networks (Shen et al. 2021) and the interactions between single IFs and MTs (Schaedel et al. 2021). In all these studies, the cytoskeletal filaments were combined in the absence of any cytolinkers.

In cells, IFs are crosslinked to F-actin and MTs via large crosslinking proteins including members of the plakin family such as plectin (Bouameur et al. 2014, Mohammed et al. 2020, Wiche and Winter 2011). Electron microscopy imaging and proximity ligation assays in cells have shown that plectin forms both IF-IF crosslinks and crosslinks of IFs to F-actin and MTs (Foisner et al. 1995, 1988, Marks et al. 2022, Svitkina et al. 1996, Wiche and Baker 1982). Vimentin filaments have been observed to closely associate with actin stress fibers (Wu et al. 2022), and there is evidence that this association depends on plectin (Jiu et al. 2015, Marks et al. 2022). Plectin has also been implicated in crosslinking of vimentin to F-actin at the base of invadopodia (Schoumacher et al. 2010, Yoneyama et al. 2014) and to the actin cortex of cells during mitosis (Serres et al. 2020). The many splice isoforms of plectin share a conserved tripartite structure with a plakin domain, central rod domain, and C-terminal domain. Depending on the isoform, plectin can have up to seven functional domains (Fig. 1A). The N-terminal motif is isoform-specific and targets plectin to specific cellular locations such as the nucleus, hemidesmosomes, or focal adhesions. Actin-binding is mediated by calponin homology domains that form an actin-binding domain (ABD) directly following the N-terminal motif (Fontao et al. 2001, Garcia-Alvarez et al. 2003). IF-binding is mediated by four domains at the C-terminus consisting of two plakin repeat domains (PRDs) separated by a linker region and a short C-terminal extremity containing Gly-Ser-Arg (GSR) repeats (Bouameur et al. 2014). The PRDs and the linker region have a similar basic groove that binds the acidic coiled-coil rod domain of IFs (Choi et al. 2002, Favre et al. 2011, Fogl et al. 2016, Kang et al. 2016, Mohammed et al. 2020).

**FIG. 1:**
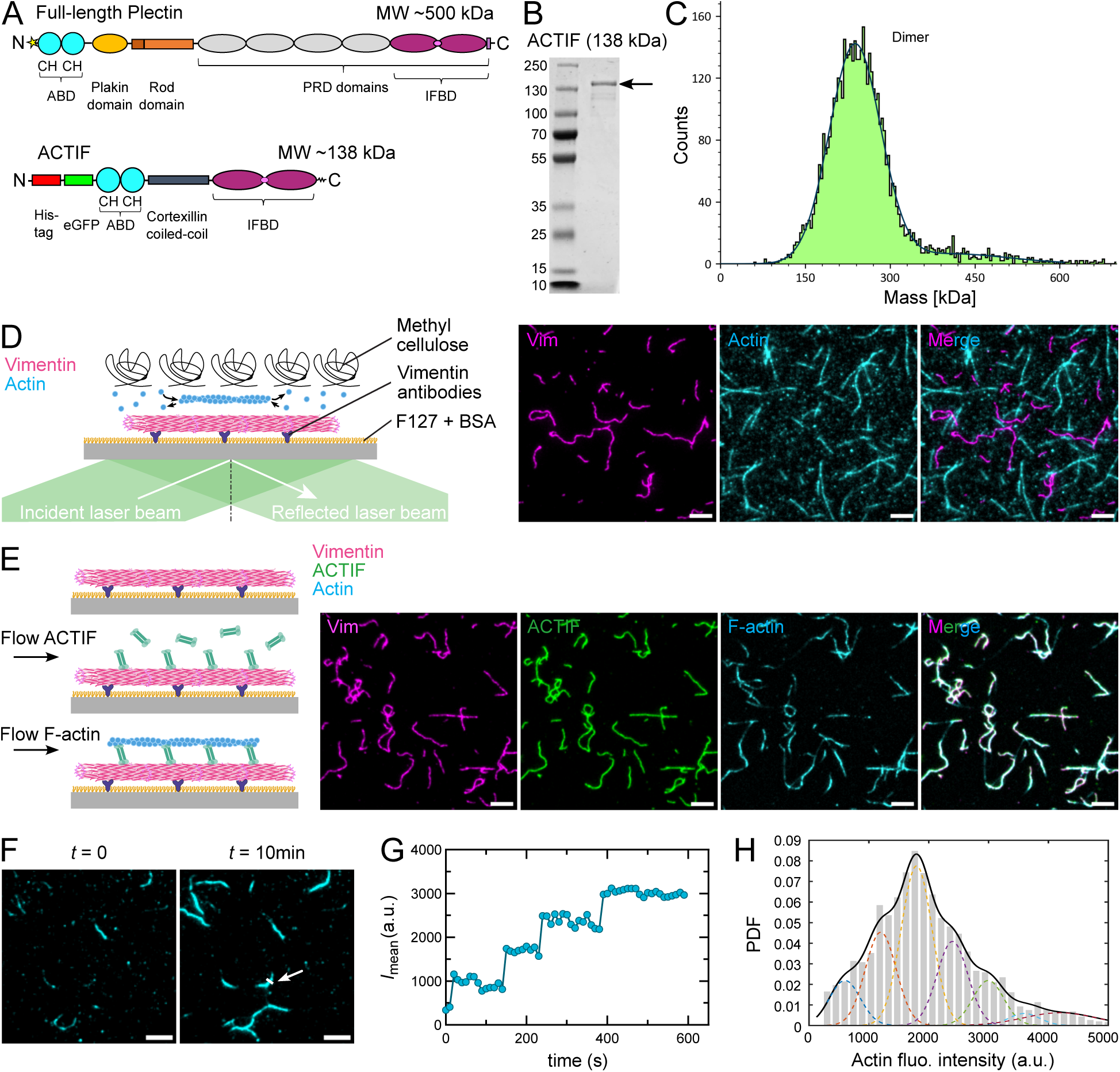
ACTIF crosslinker with the plectin IFBD binds F-actin and vimentin. A) Domain structures of full-length plectin (top) and ACTIF with the ABD from MACF and the IFBD from plectin (bottom). The star in the top schematic stands for plectin’s isoform-specific head domain. B) SDS-PAGE of ACTIF purification after gel filtration. 10 µL of proteins at 10 µM were loaded and stained with InstantBlue Coomassie Protein Stain for polyacrylamide gels. C) Interferometric scattering microscopy (iSCAT) of ACTIF preparation (50 nM in F-buffer buffer) shows a peak at a molecular weight of 243 kDa, indicating that ACTIF forms a dimer. D) Schematic of control experiment with TIRF microscopy to probe for interactions between vimentin filaments, immobilized on the microscope coverslip using vimentin antibodies, and F-actin polymerized from 1 µM G-actin in solution. Methylcellulose (0.2 %) was added to push F-actin down toward the surface. The images (right) show no co-localization between the filaments, suggesting there are no interactions. E) Schematic of sequential steps in the TIRF experiments to probe ACTIF-mediated binding of F-actin to vimentin filaments (left). First ACTIF preparation (5 nM in F-buffer) was flushed into the channel containing surface-bound vimentin filaments for 15 min. Next pre-formed F-actin (0.1 µM in F-buffer) was flushed in. Images (right) show snapshots after each step with a merged image of the final situation on the right. F) Images of F-actin attachment over time on top of ACTIF-bound vimentin filaments. G) Mean intensity over time of a representative F-actin at the cross-section marked by the white line in Fig. 1(F), showing multiple actin filaments recruited on top of ACTIF-bound vimentin. H) Distribution of pixel fluorescence intensities in the F-actin channel along vimentin filaments (~150 filaments from 2 independent experiments) at steady state. The probability distribution function (PDF) was fitted by a multiple Gaussian, with the peak positions constrained to be multiples of the position of the first peak, and common width for all the peaks except the last one. The first peak was estimated at an intensity ~600, as expected for single F-actin imaged in similar conditions. Scale bar: 5 µm.

Plectin depletion has been shown to alter the mechanical behavior of cells(Bonakdar et al. 2015, Martin et al. 2016, Moch et al. 2016, Na et al. 2009, Zrelski et al. 2023) and to impair the mechanical stability of epithelial and endothelial cell sheets (Osmanagic-Myers et al. 2015, Prechova et al. 2022). Indeed plectin mutations are associated with severe skin blistering disorders and muscular dystrophy. However, plectin’s specific role in these mechanical abnormalities as a cytoskeletal crosslinker has been difficult to decipher since it has several other roles besides direct crosslinking: it also anchors IFs to cell-matrix and cell-cell adhesion receptor complexes (Bhattacharya et al. 2009, Colburn and Jones 2018, De Pascalis et al. 2018, Geerts et al. 1999, Walko et al. 2015), to LINC complexes in the nuclear envelope (Wilhelmsen et al. 2005) and to mitochondria (Winter et al. 2008). There is hence a need for reconstitution studies where plectin’s crosslinking role can be studied in isolation from its other functions. Unfortunately, plectin is a challenging protein for reconstitution because of its giant size (500 kDa) and multidomain structure. Full-length plectin can in principle be obtained from mammalian cells (Foisner et al. 1988), but it is difficult to obtain proteins in high yield and purity using this strategy.

Here we engineer a recombinant actin-vimentin crosslinker that we call ‘ACTIF’, which is designed to crosslink F-actin and IFs in a similar manner as plectin. The engineered linker has a modular design with an N-terminal Green Fluorescent Protein (GFP) tag and actin-binding domain, a C-terminal IF-binding domain, and a coiled-coil linker to induce parallel dimerisation. Its small (138 kDa) size allows for convenient recombinant expression in bacteria. We demonstrate by imaging and rheology that ACTIF with the plectin intermediate filament binding domain (IFBD) efficiently crosslinks vimentin intermediate filaments to F-actin. Using Total Internal Reflection Fluorescence (TIRF) imaging, we quantitatively characterize the binding kinetics for the plectin-derived ACTIF crosslinker. Finally, we demonstrate the modularity of our approach by creating an alternative ACTIF variant with an IFBD based on the armadillo repeats of the tumor suppressor protein Adenomatous Polyposis Coli (APC) (Sakamoto et al. 2013). Our work provides a powerful new reconstitution platform to study the biophysical mechanisms that contribute to the functional coupling of the F-actin and IF cytoskeleton that controls cell mechanics and migration.

## 2. RESULTS

### ACTIF crosslinks F-actin and vimentin and mediates co-localization

Full-length plectin (Fig. 1A (top)) is a giant (> 500 kDa) multidomain cytolinker protein (Wiche and Winter 2011). The isoform-specific N-terminal head domain (indicated by a star in Fig. 1A, top schematic) is followed by the actin-binding domain (ABD), an elongated plakin domain (Al-Jassar et al. 2013, Ortega et al. 2016, Sonnenberg et al. 2007), a central alpha-helical rod domain (200 nm long) that mediates plectin dimerization, six tandemly arranged plakin repeat domains (PRDs, ~300 residues each) separated by linker regions of variable length, and finally, a short tail region containing Gly-Ser-Arg (GSR) repeats. The first four PRD domains do not contribute to IF-binding (Nikolic et al. 1996, Steiner-Champliaud et al. 2010). The last two PRD domains contribute together with the PRD5-PRD6 linker region (~50 residues) and the GSR-containing C-terminal extremity to ensure efficient IF binding (Bouameur et al. 2014). The linker region is essential but not sufficient for strong IF-binding (Bouameur et al. 2014, Favre et al. 2011, Nikolic et al. 1996). In addition, a sequence of approximately 50 amino acid residues (4262-4316) within one-third of the PRD5 is indispensable for filament association (Nikolic et al. 1996).

We generated a minimal crosslinker protein designed to model the crosslinking activity of plectin (Fig. 1A (bottom)), which we call ACTIF. We based the design on an earlier engineered cytolinker called TipAct that models the actin-microtubule cytolinker protein MACF/ACF7 (Alkemade et al. 2022, López et al. 2014). The TipAct protein consisted of an N-terminal green fluorescent protein (GFP) tag, followed by the F-actin binding domain of MACF, a coiled-coil spacer domain, and a C-terminal microtubule-binding domain. We retained this domain structure but replaced the microtubule-binding domain with the C-terminal intermediate filament binding domain (IFBD) of plectin. To ensure robust IF-binding, we included the last two PRD domains (PRD5 and PRD6) and their connecting linker (Nikolic et al. 1996). The ABD from MACF consists of a tandem of two calponin-homology (CH) domains and resembles the ABD of plectin (Fontao et al. 2001, Sevcik et al. 2004, Urbániková et al. 2002). The coiled-coil spacer domain separating the ABD and IFBD was taken from cortexillin I, in order to mimic plectin’s dimeric state by inducing parallel dimerization of ACTIF (Faix et al. 1996). By removing the large plakin rod and the first four PRD domains of plectin, we obtained an actin-IF cytolinker protein with a molecular weight of only 138 kDa, enabling us to express the protein in bacteria.

We first tested the purity and oligomeric state of the purified ACTIF protein. To test the protein purity, we performed SDS-PAGE analysis after the final gel filtration chromatography step (see Supplemental Figure 2). As shown in Fig. 1B, the SDS-PAGE gel shows a major band localized at the expected molecular weight of 138 kDa and smaller bands indicative of protein degradation. Densitometry analysis showed that full length ACTIF made up 64% of the protein preparation (Supplemental Figure 3). To test whether the smaller molecular weight products are degradation products of ACTIF, we tested for the presence of the N-terminal eGFP tag in the bands by fluorescence imaging of the gels using a Typhoon imager (Supplemental Figure 4) and by Western blot analysis using antibodies to detect GFP (Supplemental Figure 5). Both experiments showed that all the proteins present in the ACTIF preparation contained eGFP, indicating that the lower molecular weight products are indeed degradation products of ACTIF. Finally, we performed liquid chromatography–mass spectrometry (LC–MS) analysis of the ACTIF preparation (Supplemental Figure 6). This analysis confirmed that the most abundant protein in the sample had a molecular weight compatible with the expected value for ACTIF at 138,747 Da, with a precision of about 1 Da. Putative identification of some of the mass peaks allowed us to identify ACTIF fragments, consistent with the conclusion from Typhoon and Western blot analysis that these proteins result from partial proteolysis of ACTIF. To check the oligomeric state of the ACTIF protein in solution, we performed interferometric scattering (iSCAT) microscopy (Young et al. 2018). With iSCAT the light scattering of molecules near a surface is amplified with reference light, typically the reflection of the laser on the bottom of the glass slide (Supplemental Figure 1). Background subtraction makes it possible to detect the molecules landing on or moving away from the surface. The detected signal is directly proportional to the molecular weight of the proteins (Young et al. 2018). As shown in Fig. 1C), the mass histogram confirms that the crosslinker is dimeric, as shown by the peak of 98% of signal counts being centered around 243 ± 73 kDa, about twice the molecular weight of the monomer. Note that this peak likely includes dimers of full length ACTIF as well as dimers of the degradation products.

Before testing the crosslinking activity of ACTIF, we first wanted to test whether F-actin and vimentin have any direct interactions under our experimental conditions. This idea was motivated by earlier findings that vimentin can bind F-actin with its C-terminal tails (Esue et al. 2006). To test for interactions, we used a TIRF microscopy assay (Fig. 1D (left)). We prepared a microfluidic flow chamber made of coverglass treated with vimentin antibodies that securely trap vimentin filaments, and with passivation agents including pluronic F127 and bovine serum albumin (BSA), to prevent unwanted protein binding to the substrate. Pre-assembled vimentin filaments fluorescently labeled with Alexa Fluor 555 were next introduced into the chamber and allowed to adhere to the substrate. Next, we flowed in ATTO-647-labeled G-actin monomers in F-buffer supplemented with 0.2 % methylcellulose, to polymerize F-actin directly in the chamber. Methylcellulose was used to force the growing F-actin filaments close to the substrate with vimentin. As shown in Fig. 1D (right), we did not observe any signs of co-localization between F-actin and the fixed vimentin filaments from the beginning of the F-actin polymerization until they became long filaments. Timelapse movies confirmed that the F-actin filaments are freely moving (Supporting Movie 1). We conclude that vimentin and F-actin do not interact directly under the conditions of our assay. Moreover, since G-actin co-existed with F-actin during the *in situ* assembly, the absence of colocalization of actin and vimentin signals suggested that neither G-actin nor F-actin bind vimentin filaments in our experimental conditions, although we cannot rule out that transient weak interaction may take place.

Having established that vimentin and F-actin do not significantly interact, we could then test whether ACTIF induces interactions. We again used a TIRF imaging assay where we sequentially flushed in the different components (Fig. 1E (left)). We first flushed in 37 nM of pre-assembled vimentin and allowed these filaments to adhere to the vimentin antibodies on the substrate. We previously showed that vimentin attachment to a glass surface through antibodies has minor impact on the structural organization of the filaments Nunes Vicente et al. (2022). When we next flushed in 5.3 nM of ACTIF, we observed that it rapidly bound to the vimentin filaments as seen by co-localization (Fig. 1E (middle)). Finally, we flushed in 0.1 µM of pre-assembled F-actin. Note that as the F-actin was diluted 100-fold just before use from a stock solution of 10 µM F-actin pre-polymerized for 1 h, we assumed the concentration of G-actin to be in the nM range (100 times smaller than the critical concentration for actin polymerization), thus providing minimal contribution to the fluorescent signal in the actin channel. Over time, we observed that the actin filaments settled and remained on top of the vimentin filaments (Fig. 1E (right)). At steady state, F-actin, vimentin, and ACTIF co-localized, as shown in the merged image (Fig. 1E (far right) and Supporting Movie 2). Following the maximum intensity of the F-actin signal along a cross-section drawn perpendicularly to the vimentin filaments, we could follow the time evolution of successive F-actin binding events, which appeared as intensity steps (Fig. 1G). By quantifying pixel fluorescence intensities in the F-actin channel along vimentin filaments at steady states (~ 150 filaments from 2 independent experiments), we observed a complex distribution with multiple peaks indicating the presence of multiple actin filaments (Fig. 1H), different from the single peak distribution obtained with vimentin only (Supplemental Figure 7). As the intensity of the first peak matched the intensity expected for single F-actin, we estimated that an average of 3 actin filaments and up to 7 actin filaments could bind a vimentin filament through ACTIF crosslinking at steady state (Fig. 1H). These observations prove that ACTIF is capable of binding both F-actin and vimentin and of crosslinking the two filament types.

### ACTIF robustly binds and couples vimentin and F-actin filaments

We performed a series of TIRF microscopy assays to validate the efficiency of ACTIF in crosslinking actin and vimentin filaments in real-time. In the similar TIRF microscopy and flow chamber setup as shown in Fig. 1(D and E), we flowed ACTIF at a low concentration of *c* ≈ 5 nM into a channel with surface-bound vimentin filaments (Fig. 2A). We observed rapid co-localization of ACTIF with vimentin within 10 min (Fig. 2B). Using the coloc 2 plugin from Fiji, we quantified the colocalization between the different components and obtained Manders coefficients of *M* 1 = 0.978 and *M* 2 = 0.984 for the colocalization of ACTIF vs. vimentin and *M* 1 = 0.972 and *M* 2 = 0.987 for the colocalization of F-actin and vimentin. Fluorescence quantification showed that ACTIF bound to vimentin with a reaction rate *K* ≈ 0.0140 ± 0.0024 s^−1^ (mean ± SD, averaged over 3 independent repeats) obtained by fitting the kinetics of association with a single exponential (Fig. 2D). Once the ACTIF signal reached a steady state (after 10 minutes), we performed a Fluorescence Recovery After Photobleaching (FRAP) assay (Fig. 2A) to measure the ACTIF unbinding rate, assuming that the recovery is reaction-limited (Bulinski et al. 2001). As shown in Fig. 2C and quantified in Fig. 2E, we observed about 30 % of ACTIF is mobile (*N* = 3 independent repeats). We presume that the mobile fraction corresponds to dimeric ACTIF and the immobile fraction to higher oligomerization of ACTIF induced by attachment to vimentin, which increases protein avidity. By fitting the recovery curve to a single exponential, we estimated the unbinding rate of the mobile fraction of ACTIF from vimentin filaments of *k_off_* ≈ 0.0032 ± 0.0012 s^−1^ (mean ± SD, *N* = 3 repeats). Moreover, according to densitometry and Typhoon assays (Supplemental Figure 3 and 4), only 64 % of the ACTIF preparation was full-length with a functional IFBD domain, which reduced the binding efficiency of ACTIF on vimentin filaments. With the assumption of a first-order reaction with binding efficiency of 65 % and assuming that 30 % of ACTIF is mobile, the rate *K* of association kinetics can be approximated by: *K* = 0.65 *k*_on_ *c* + *f*_m_ *k*_off_, where *k*_on_ is the binding rate and *f*_m_ is the mobile fraction. From the concentration *c*, the rate *K* and the unbinding rate *k*_off_ estimated by FRAP, we estimated the binding rate of ACTIF to vimentin *k*_on_ ≈ (3.7 ± 0.7) · 10^6^ M^−1^s^−1^ (mean ± SD, *N* = 3 repeats), and the affinity described by a dissociation equilibrium constant *K_d_* ≈ 0.9 ± 0.4 nM (*N* = 3 repeats). We next tested the F-actin crosslinking ability of the vimentin-bound ACTIF by flowing in pre-assembled F-actin in a buffer without methylcellulose. We observed that the F-actin filaments quickly attached and localized along vimentin filaments, forming composites of F-actin-ACTIF-vimentin (Fig. 2F). Interestingly, once the composites were established, ACTIF was fixed inside the composites as its fluorescence intensity remained constant over time when we flushed in a solution that contained F-actin but no ACTIF (Fig. 2G, green curve). We verified that ACTIF partially dissociated from vimentin in the absence of F-actin when the ACTIF soluble pool was flushed out (Fig. 2G, orange curve), indicating that ACTIF is stabilized on vimentin through its binding to actin. Note that this experiment also demonstrates that the incomplete fluorescence recovery seen in FRAP experiments is not due to any protein alteration by bleaching. Our findings suggest strong binding of ACTIF to vimentin filaments and stable crosslinking between vimentin and F-actin filaments.

**FIG. 2:**
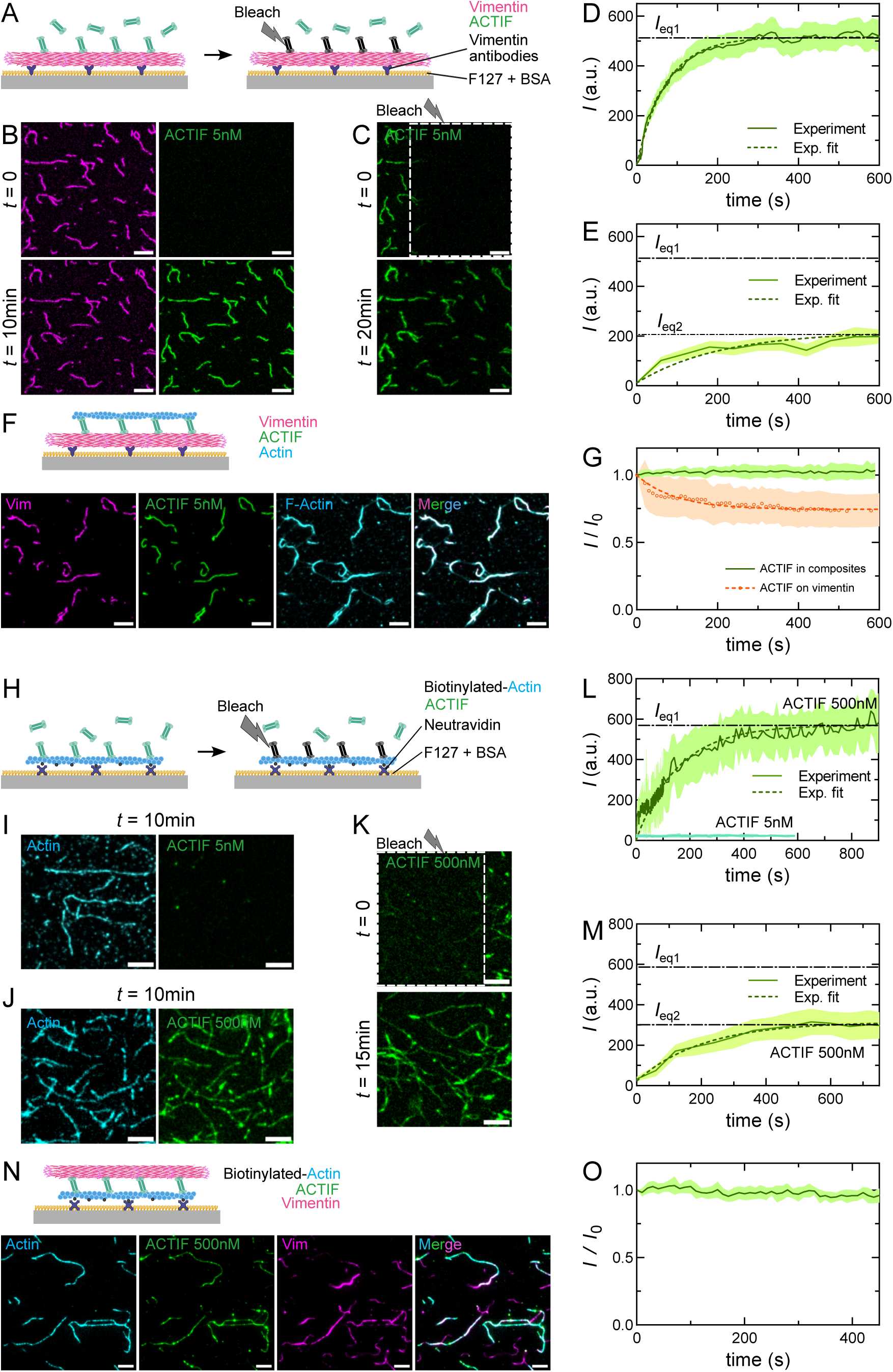
TIRF microscopy experiment to elucidate the crosslinking efficiency of ACTIF. A) Assay to measure the binding rate of ACTIF to surface-bound vimentin by imaging (left) and the unbinding rate by FRAP (right). B) Still images at *t* = 0, directly after flushing in 5 nM ACTIF preparation, and at *t* = 10 min, showing ACTIF (green) co-localizing with vimentin filaments (magenta). C) Images showing the recovery of ACTIF fluorescence at *t* = 20 min after photobleaching the region boxed by white dashed lines at *t* = 0. D) Mean fluorescence intensity of ACTIF on vimentin filaments (averaged over 30 filaments) over time, obtained from the representative repeat shown in Fig. 2(B). The data is fitted by an exponential equation, *I* = *I*_eq1_ (1 − *e^−Kt^*). E) Recovery of fluorescence intensity of ACTIF on vimentin filaments (mean fluorescence averaged over 30 filaments) over time after photobleaching, obtained from the representative repeat shown in Fig. 2(C). The data is fitted by an exponential equation, *I* = *I*_eq2_ (1− *e*^−*k_off_t*^). F) Assay and images of steady-state vimentin-ACTIF-F-actin composites with surface-bound vimentin. G) Normalized fluorescence intensity of ACTIF in the actin-vimentin composites (green) over time after flushing in a solution containing F-actin but no ACTIF (at *t* = 0), and on vimentin (orange) over time after flushing out solution-phase ACTIF (averaged over 30 and 50 filaments respectively). H) Assay to measure the binding rate of ACTIF to surface-bound F-actin by imaging (left) and the unbinding rate by FRAP (right). I, J) Images of ACTIF (green) co-localizing with actin filaments (cyan) at *t* = 10 min after flushing in ACTIF at a concentration of ~5 nM (I) and ~500 nM (J). K) Images showing the recovery of ACTIF fluorescence at 500 nM on F-actin at *t* = 15 min after photobleaching the region boxed by white dashed lines at *t* = 0. L) Fluorescence intensity of ACTIF preparation at concentrations of 5 nM and 500 nM on F-actin (both conditions averaged over 30 filaments) over time, obtained from the representative repeat shown in Fig. 2(J). The data is fitted by an exponential equation, *I* = *I*_eq1_ (1 − *e^−Kt^*). M) Recovery of fluorescence intensity of ACTIF preparation at 500 nM on F-actin (averaged over 30 filaments) over time after photobleaching, obtained from the representative repeat shown in Fig. 2(K). The data is fitted by an exponential equation, *I* = *I*_eq2_ (1 − *e*^−*k_off_t*^). N) Assay and images of steady-state actin-ACTIF-vimentin composites with surface-bound F-actin. O) Normalized fluorescence intensity of ACTIF in the composites over time, after flushing in F-actin while flushing out solution-phase ACTIF at *t* = 0 (averaged over 30 filaments). Shaded areas in all the graphs show the standard deviations over 30 to 50 filaments. Dashed lines show exponential fits for the kinetics of association and for FRAP recovery curves. Scale bars: 5 µm.

To study the binding and unbinding rates of ACTIF on F-actin, we modified the experimental setup by immobilizing pre-assembled biotinylated actin filaments on the surface of the flow chamber, which had been coated with neutravidin, and the passivation agents F127 and BSA, respectively. Next, we flowed in ACTIF to characterize the kinetics of association to F-actin, then performed FRAP to measure the unbinding rate from F-actin and deduce the binding rate to F-actin, and finally flowed in vimentin filaments to create F-actin-ACTIF-vimentin composites (Fig. 2(H, N)). When we flowed in ACTIF at the same concentration of ~5 nM, as in the earlier experiment with immobilized vimentin, we did not observe any signs of binding to the immobilized F-actin (Fig. 2(I, L) after 10 minutes. To our surprise, we needed to flow in *c* ≈ 500 nM of ACTIF to achieve a final ACTIF fluorescence intensity equivalent to its intensity on vimentin achieved with a 100-fold lower ACTIF concentration (Fig. 2J). Although it is expected that the number of binding sites for ACTIF on F-actin is not the same as that on vimentin (see Discussion), the difference by a factor of 100 in concentration indicates the binding rate is much lower on actin than on vimentin. By fitting the time-dependent increase of the ACTIF fluorescence intensity to a single exponential, we obtained a reaction rate *K* ≈ 0.0080 ± 0.0013 s^−1^ (mean ± SD, *N* = 3 repeats) (Fig. 2L). FRAP measurements on the actin-bound ACTIF allowed us to estimate the mobile ACTIF unbinding rate of *k*_off_ ≈ 0.0064 ± 0.0011 s^−1^ from F-actin, and a mobile fraction of about 48 % (*N* = 3) (Fig. 2K, M). The binding efficiency of ACTIF on F-actin is not affected by the 65 % purity of the protein since the ABD domain on the N-terminus is always intact (See Supplemental Figure 4). Hence, the association kinetics of ACTIF on F-actin is estimated by *K* = *k*_on_ *c* + *f*_m_ *k*_off_. From the values of *K*, *c*, *k*_off_ and *f*_m_, we estimated the binding rate to F-actin *k*_on_ ≈ (9.5±2.9)·10^3^ M^−1^s^−1^ and the affinity to F-actin described by *K_d_* ≈ 705 ± 137 nM. When we finally flushed in pre-assembled vimentin filaments, we again obtained colocalization of the vimentin filaments with the ACTIF-saturated F-actin, reaffirming the robust formation of the composite (Fig. 2N). Time-lapse imaging also again showed the ACTIF signal in the composite structure to be constant, even though no ACTIF was supplied in the solution (Fig. 2O). Overall, our results indicate the binding of ACTIF to F-actin is about 500 times lower than its affinity to vimentin (summary in Table 1).

**TABLE 1:** Summary of (un)binding rate constants and affinities of ACTIF towards vimentin and actin filaments measured by TIRF imaging and FRAP assays. Values shown in Repeat 1-3 are means averaged over 30 filaments calculated for each independent repeat experiment. The combined “Mean ± SD” represents the averaged mean from 3 repeats with SD being the standard deviation of the means.

| | Repeat # | $K$ ($\text{s}^{-1}$) | $k_{\text{on}}$ ($\text{M}^{-1} \text{s}^{-1}$) | % mobile<br>$f_{\text{m}}$ | Mobile fraction | |
| --- | --- | --- | --- | --- | --- | --- |
| | | | | | $k_{\text{off}}$ ($\text{s}^{-1}$) | $K_{\text{d}}$ (nM) |
| ACTIF<br>on<br>vimentin<br>$c = 5.3$ nM | 1 | 0.0111 | $3.1 \cdot 10^6$ | 25 % | 0.0023 | 0.75 |
| | 2 | 0.0170 | $4.7 \cdot 10^6$ | 27 % | 0.0025 | 0.53 |
| | 3 | 0.0138 | $3.4 \cdot 10^6$ | 40 % | 0.0049 | 1.43 |
| | Mean ± SD | $0.0140 \pm 0.0024$ | $(3.7 \pm 0.7) \cdot 10^6$ | $(31 \pm 7) \%$ | $0.0032 \pm 0.0012$ | $0.9 \pm 0.4$ |
| ACTIF<br>on<br>F-actin<br>$c = 530$ nM | 1 | 0.0079 | $7.7 \cdot 10^3$ | 56 % | 0.0068 | 883 |
| | 2 | 0.0096 | $13.6 \cdot 10^3$ | 32 % | 0.0075 | 551 |
| | 3 | 0.0065 | $7.2 \cdot 10^3$ | 55 % | 0.0049 | 681 |
| | Mean ± SD | $0.0080 \pm 0.0013$ | $(9.5 \pm 2.9) \cdot 10^3$ | $(48 \pm 11) \%$ | $0.0064 \pm 0.0011$ | $705 \pm 137$ |

### The plectin IFBD induces vimentin bundling

To test whether the isolated plectin IFBD domain is capable of binding or bundling vimentin, we expressed and purified this domain, comprising PRD 5 and PRD 6 and the intervening linker region that provides the major IF-binding site of plectin (Nikolic et al. 1996). iSCAT imaging showed the isolated IFBD to form mostly monomers with a small (13 %) fraction of dimers (Supplemental Figure 8A) and SDS-PAGE analysis showed a major band at the expected molecular weight of 71 kDa (Supplemental Figure 8B). Next, we co-polymerized the IFBD with vimentin filaments at different IFBD:vimentin molar ratios, always keeping the vimentin molar concentration fixed at 4 µM, and imaged the mixtures by electron microscopy (EM). EM provides higher spatial resolution than TIRF imaging and allows us to resolve filament and bundle morphologies and widths. Vimentin filaments reconstituted in the absence of the IFBD were visible in EM images as semiflexible filaments (Fig. 3A (left)). To determine the filament widths, we plotted intensity profiles perpendicular to the segment of interest at multiple locations along the filament contours. We then measured the width of the bright area between the darker borders, which corresponds to the filament diameter. We observed an average filament width of ~10 nm (Supplemental Figure 9), in agreement with previous literature (Wen and Janmey 2011). Surprisingly, when vimentin was co-polymerized with the IFBD at a molar ratio of IFBD:vimentin of 1:10 (0.4 µM of IFBD), it formed thin bundles (see Fig. 3A (middle)). Bundles often contained around three to four distinguishable filaments. Using the same procedure as described above to determine the bundle widths, we found that the bundles had a variable width with an average bundle size of 33 ± 12 nm (Fig. 3B). Upon increasing the IFBD:vimentin molar ratio to 1:1 (4 µM of IFBD), vimentin filaments formed straight and well-aligned bundles (Fig. 3A (right)) of an average diameter of 58 ± 22 nm (Fig. 3B). The bundles formed at a 1:1 IFBD:vimentin ratio were significantly thicker than those formed at a 1:10 ratio (*p <* 0.0001, calculated by a statistical unpaired t-test). We conclude that the IFBD domain can by itself bundle vimentin filaments at high (> 400 nM) concentrations. Presumably, bundling requires dimerization or clustering of the IFBD domain, which we showed to be mostly monomeric (with only 13 % dimeric) in iSCAT measurements performed at a concentration of 40 nM (Supplemental Figure 8A). Potentially, at the high (> 400 nM) concentrations used in the EM assays, the IFBD domain can oligomerize even more. As a control, we also tested co-polymerized actin and IFBD. In this case, EM images showed no signs of actin bundling, but plectin was present in the form of dense globular structures, suggesting that the IFBD can cluster at high concentrations (Supplemental Figure 11).

**FIG. 3:**
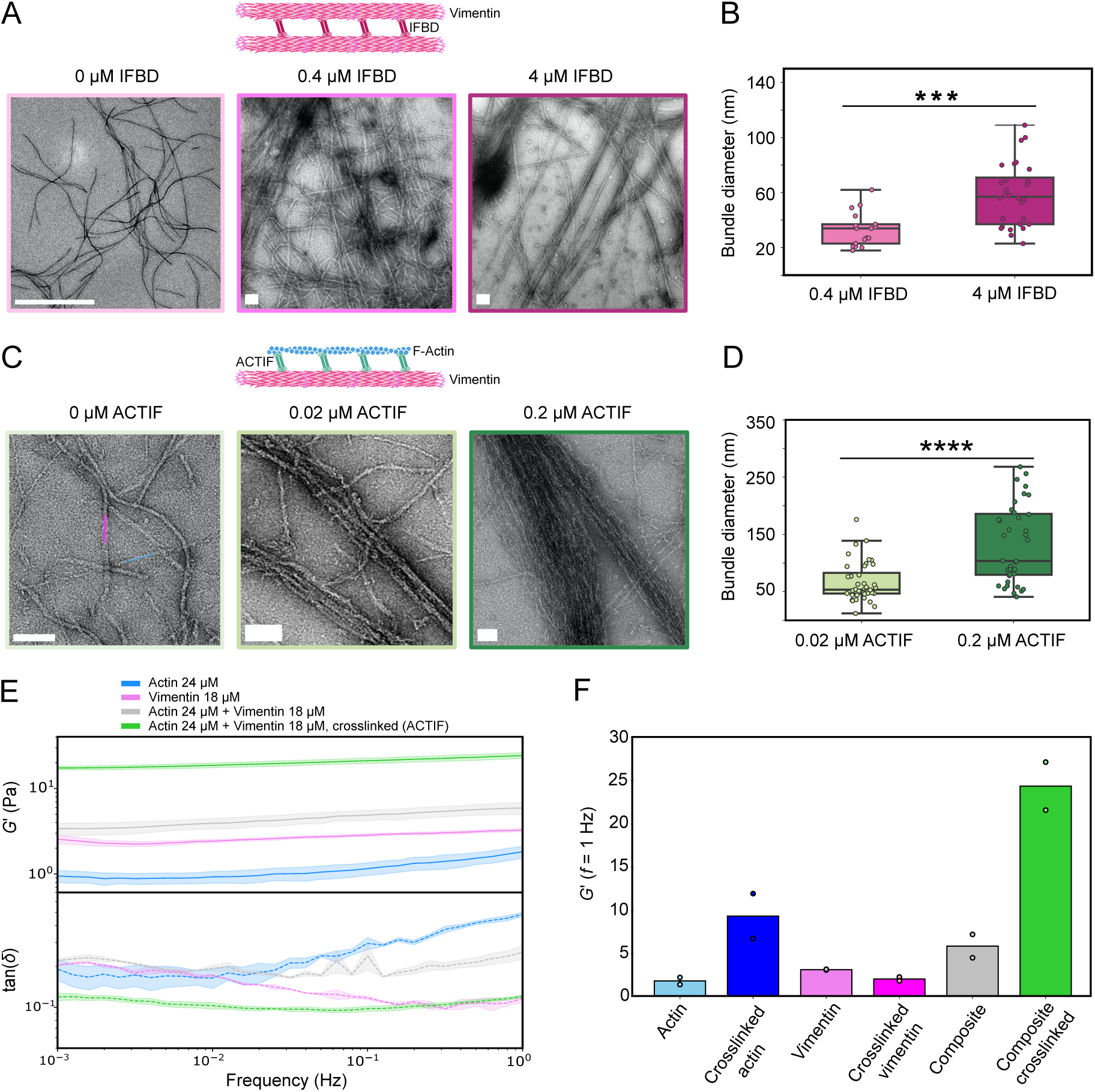
Electron microscopy (EM) and rheology experiments to test binding and crosslinking activities of ACTIF preparation and its plectin IFBD. A) EM images of reconstituted vimentin filaments alone (left) or co-polymerized with 0.4 µM (middle) or 4 µM (right) of IFBD (schematic on top). Enlarged versions are shown in Supplemental Figure 10). Corresponding box plots of vimentin bundle widths, with 17 bundles measured for 0.4 µM of IFBD and 28 bundles measured for 4 µM of IFBD. Two independent sample replicates per condition were imaged. *** *p <* 0.001 from unpaired *t*-test. C) EM images of co-polymerized F-actin (cyan) and vimentin filaments (pink) without ACTIF preparation (left) or with 0.02 µM (middle) or 0.2 µM (right) of ACTIF preparation (schematic on top). D) Corresponding box plots of the F-actin/vimentin co-bundles. **** *p <* 0.0001 from unpaired *t*-test. For both conditions, 40 bundles were measured. Two independent sample replicates per condition were imaged. E) (Top) Frequency dependence of the storage modulus of crosslinked (green) and non-crosslinked (grey) actin/vimentin composite networks and of one-component actin (blue) or vimentin (pink) networks. Protein concentrations: 24 µM F-actin, 18 µM vimentin. (Bottom) Corresponding loss tangents, defined as the ratio of the loss modulus *G′′* to the storage modulus *G′* (same color code). Note that the loss tangent is noisy because the applied strain and the shear moduli (especially *G′′*) are small, so the stress is close to the sensitivity limit of the rheometer. F) Bar plots comparing the elastic modulus *G′* at a frequency of 1 Hz of entangled and crosslinked networks of actin, vimentin, and actin/vimentin composites. Protein concentrations: 24 µM F-actin, 18 µM vimentin, 4.2 µM ACTIF preparation. For each condition, *N* = 2, circles are individual data points. All scale bars are 100 nm, except in the far left image of panel A, where the scale bar is 1000 nm.

### ACTIF forms bundles of F-actin and vimentin filaments

With the certainty based on EM data that the IFBD from plectin can bind vimentin filaments, we next imaged mixtures of F-actin and vimentin filaments with and without ACTIF, always keeping the monomer concentrations at 1 µM for actin and 1µM for vimentin. In the absence of ACTIF, actin filaments can be distinguished from vimentin filaments by their smaller diameter (8 nm vs 10 nm) (Wen and Janmey 2011)) and by their larger persistence length (10 µm vs 1 µm (Wen and Janmey 2011)). As shown in Figure 3 C(left) and Supplemental Figure 12, the two filament types can indeed be easily distinguished. Upon adding ACTIF in a molar ratio of 1:100 to the sum of the vimentin and actin molar concentrations, we found that tightly packed bundles formed (Fig. 3C (middle)) with an average diameter of 66 ± 34 nm (Fig. 3D). Note that bundling of actin and vimentin by ACTIF occurs at a much lower crosslinker:monomer ratio than bundling of vimentin by the IFBD domain, consistent with ACTIF being a parallel dimer presenting two actin-binding domains on one end and two IF-binding domains on the other end. At a ten-fold larger ACTIF:monomer ratio of 1:10 (0.2 µM of ACTIF), even thicker bundles were observed that consisted of a larger number of filaments (Fig. 3C (right)). The average bundle size was significantly larger (131 ± 66 nm) than for the bundles formed at the 1:100 molar ratio (Fig. 3D), *p <* 0.00001). Interestingly, the bundle widths showed an apparently bimodal distribution at the 1:10 ACTIF:monomer ratio, with the first peak centered around a similar width of 70 nm as observed at 1:100. This observation suggests a hierarchical bundling process, where thinner bundles can get connected into thicker bundles. Many bundles were too dense to observe their constituent filaments, but in some cases we could observe bundles clearly consisting of a mixture of actin and vimentin filaments (Supplemental Figure 13A), some bundles eventually splitting into two or more sub-bundles (Supplemental Figure 13B) and some bundles apparently consisting only of actin filaments (Supplemental Figure 13C). Note that, the presence of 36 % of truncated ACTIF that does not bind vimentin may impact the morphology and composition of actin bundles. We conclude that ACTIF drives the formation of mixed actin-vimentin bundles in a concentration-dependent manner.

### Composite F-actin/vimentin networks stiffen upon crosslinking with ACTIF

To test whether ACTIF is capable of crosslinking F-actin and vimentin in dense networks, we used shear rheology to measure the viscoelastic properties of composite networks formed with or without ACTIF. The viscoelastic response of cytoskeletal networks is known to be sensitive to the presence of crosslinkers, which tend to enhance the elastic modulus and suppress viscous dissipation (Ward et al. 2008). We polymerized actin (24 µM), vimentin (18 µM), or composite actin-vimentin networks between the plates of a rheometer. We did not observe any major differences in the polymerization kinetics between one-component versus two-component networks (Supplemental Figure 14). After 2 hours, when network formation was completed, we measured the linear viscoelastic response by applying small amplitude oscillatory shear with a strain amplitude of 1% while varying the frequency logarithmically between 0.001 and 10 Hz. We determined the network elastic (storage) shear modulus, *G′*, and viscous (loss) modulus, *G′′*, from the stress response. In the absence of ACTIF, the composite networks had a small elastic modulus of order ~ 10 Pa that increased very weakly as an approximate power law with frequency with an exponent of 0.07 (grey curve in Fig. 3E). In the presence of ACTIF (4.2 µM), the networks were more than an order of magnitude stiffer (green curve in Fig. 3E), as summarized in Fig. 3F. The frequency dependence for the crosslinked network again followed a weak power law with an exponent of 0.07. Simultaneously, the network with ACTIF was more solid-like, as characterized by a smaller loss tangent, 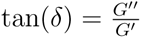, compared to the corresponding network without ACTIF (Fig. 3F)). The rheology measurements hence prove that ACTIF crosslinks the composite network.

To test whether there is mechanical synergy upon combining actin and vimentin, we also performed rheology measurements for isolated actin networks (blue curve in Fig. 3E) and vimentin networks (pink curve in Fig. 3E). The stiffness of the actin/vimentin composite network formed without ACTIF was approximately equal to the sum of the elastic moduli of the two isolated networks (Fig. 3F). Interestingly, when actin was polymerized in the presence of ACTIF (4.2 µM), it formed stiffer networks than in the absence of ACTIF (Supplemental Figure 15A), indicating that ACTIF is capable of crosslinking actin filaments. By contrast, when vimentin was polymerized in the presence of ACTIF, it formed a slightly softer network than in the absence of ACTIF (Supplemental Figure 15B). This suggests that ACTIF may partially disrupt crosslinking by vimentin’s C-terminal tail domains (Aufderhorst-Roberts and Koenderink 2019) or that ACTIF generates network inhomogeneities by bundling vimentin filaments. Interestingly, the stiffness of the crosslinked actin/vimentin composite network is larger than the sum of the moduli of the two isolated networks (Fig. 3F), suggesting the presence of some form of mechanical synergy.

### ACTIF crosslinker variant with the IFBD of APC

The engineered ACTIF crosslinker has a modular design such that its functional modules can be swapped for alternative domains. To prove the benefit of this modularity, we swapped the IFBD from plectin for an alternative IFBD, taken from the Adenomatous Polyposis Coli (APC) protein. APC was reported to directly bind vimentin filaments in astrocytes, leading to crosslinking and elongation of vimentin filaments along microtubules (Sakamoto et al. 2013). IF-binding was traced to the N-terminal Armadillo repeats of APC via biochemical assays with truncated APC proteins. We hence engineered an ACTIF variant where the IFBD from plectin was substituted by the armadillo repeats of APC, specifically, the region referred to as APCn2 (Sakamoto et al. 2013). The ACTIF-APCn2 crosslinker therefore consisted of an N-terminal GFP-tag, followed by the F-actin binding domain of MACF2, the cortexillin I coiled-coil to induce dimerization, and finally a C-terminal region containing the armadillo repeats of APC (Fig. 4A). We performed iSCAT to check the oligomeric state of the ACTIF-APCn2 crosslinker and found it to be dimeric (Supplemental Figure 8C). To test the protein purity, we performed SDS-PAGE analysis after the final gel filtration chromatography step. As shown in Supplemental Figure 8D, the SDS-PAGE gel shows a major band localized at the expected molecular weight of 116 kDa and a smaller second band at ~80 kDa indicative of some protein degradation.

**FIG. 4:**
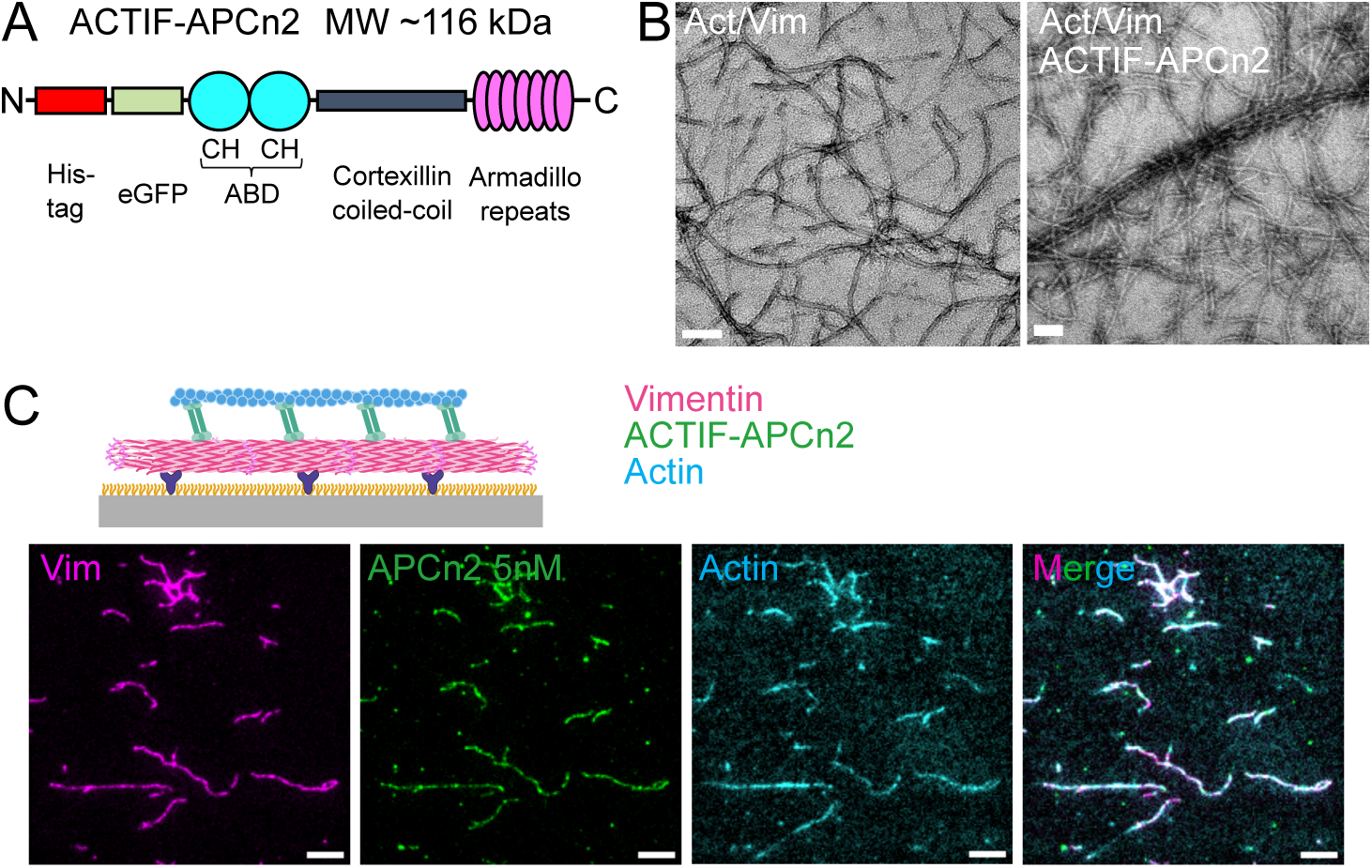
ACTIF crosslinker with the IFBD of APC binds and crosslinks F-actin and vimentin. A) Schematic representation of the engineered crosslinker ACTIF-APCn2, which is identical to ACTIF aside from its IFBD domain consisting of the armadillo repeats of APC. B) EM image of actin 1.5 µM and vimentin 1.5 µM co-polymerized in the absence (left) and presence (right) of 0.3 µM ACTIF-APCn2. ACTIF-APCn2 forms thick straight bundles. Scale bars: 100 nm. C) Schematic and fluorescent images from the TIRF microscopy assay following the same protocol as Fig. 1(E), showing F-actin was added in solution and recruited on top of ACTIF-APCn2-bound vimentin after ACTIF-APCn2 at ~5 nM was flushed into the chamber with vimentin filaments on the substrate. Scale bars: 5 µm.

We first performed EM assays to test the ability of ACTIF-APCn2 to crosslink F-actin and vimentin filaments. We compared actin and vimentin mixtures co-polymerized in the absence (Fig. 4B (left)) and presence (Fig. 4B (right)) of 0.3 µM of ACTIF-APCn2. ACTIF-APCn2 caused the formation of thick straight bundles of F-actin and vimentin filaments, showing that it is capable of crosslinking the two filament types. ACTIF-APCn2 induces bundles that are strikingly long (Supplemental Figure 16A) and that are thinner than ACTIF-induced bundles (average width of 58 ± 8 nm at 0.3 µM of ACTIF-APCn2, Supplemental Figure 16B). We next performed TIRF microscopy to validate the efficacy of ACTIF-APCn2 in crosslinking actin and vimentin filaments in real-time. After immobilizing vimentin filaments on a substrate as described earlier, we sequentially flushed in ACTIF- APCn2 (~5 nM) for 10 min, then pre-assembled F-actin. We observed binding of ACTIF-APCn2 to vimentin and progressive recruitment of F-actin on top of the ACTIF-APC2-decorated vimentin filaments (Fig. 4C), forming a stable actin-ACTIF-APCn2-vimentin composite. Quantative colocalization analysis gave Manders coefficients of *M* 1 = 0.932 and *M* 2 = 0.929 for the colocalization of ACTIF-APCn2 on vimentin and *M* 1 = 0.906 and *M* 2 = 0.916 for the colocalization of actin on vimentin, similar to the values observed with ACTIF. When we characterized the ACTIF-APCn2 association on vimentin at a similar concentration as for ACTIF (5 nM), we noted that ACTIF-APCn2 associated at a rate about 60 times lower than ACTIF during the initial phase of fluorescence intensity increase (Supplemental Figure 17). We conclude that the type of IFBD impacts the association rate of the ACTIF on vimentin.

## 3. DISCUSSION

In this work, we reconstituted specific interactions between actin and vimentin filaments by engineering a cytolinker protein that recapitulates the crosslinking activity of plectin. We found that the engineered cytolinker protein, which we named ACTIF, has a high binding affinity (*K*_d_ ≈ 1.4 nM) for vimentin and lower binding affinity for actin (*K*_d_ ≈ 0.7 µM). TIRF imaging showed that ACTIF molecules very stably crosslink actin and vimentin filaments since we did not observe any dissociation when we removed the soluble ACTIF pool, while ACTIF partially dissociated when only attached to vimentin. ACTIF is thus able to form long-lived composite actin-vimentin bundles.

The affinity of ACTIF on vimentin and on actin was estimated from the mobile fraction of ACTIF in FRAP experiments, which we believe is dimeric (~30% on vimentin and ~50% on actin). We presume that the high proportion of immobile ACTIF results from oligomerization induced by the high density of ACTIF on the filaments, which increases the strength of the bond between the oligomers and the filaments. This could explain why we only observed one population of dimers of ACTIF with the iSCAT experiments in similar conditions (concentration and buffer), but two populations on filaments (dimers and oligomers induced by the high density on filaments). We believe that the ACTIF immobile fraction is more important on vimentin than on F-actin due to the higher density of binding sites on vimentin (at least 800 per micron length of vimentin filaments made of 10 tetramers per cross-section (Eibauer et al. 2023) and an axial repeat of 50 nm (Nunes Vicente et al. 2022), compared to 360 per micron length of F-actin (Bremer et al. 1991) if we assume one ACTIF per monomer of vimentin or actin).

The high affinity of ACTIF for vimentin is qualitatively consistent with previous biochemical measurements using different plectin fragments, showing that the PRD5 and PRD6 domains and intervening linker constitute the main vimentin binding site. In quantitative terms, ACTIF under the conditions of our assays had a higher binding affinity for vimentin as compared to values reported previously for these fragments, specifically fragments consisting of the PRD5 domain and linker (*K*_d_ = 50 nM) (Nikolic et al. 1996), the PRD5-PRD6 (including the linker region) (*K*_d_ = 0.15 − 0.3 µM) (Spurny et al. 2007), or the entire C-terminus starting from the PRD5 domain (*K*_d_ = 135 nM) (Bouameur et al. 2014). Variants of ACTIF with IFBDs composed of different combinations of domains could be used in the future to sort out the contribution and synergy of the different domains of plectin implicated in controlling its affinity for vimentin. Also, ACTIF variants could be used for testing the binding affinity and crosslinking efficiency of PRD and linker domains from other plakin proteins. Sequence analysis has shown that these domains differ in the number of basic residues in their IF-binding groove, suggesting that they present a spectrum from weak to strong binding affinities (Mohammed et al. 2020). The lower binding affinity of ACTIF’s ABD for F-actin is consistent with previous measurements for TipAct (López et al. 2014), the engineered actin-microtubule crosslinker that formed the template for ACTIF. ACTIF has the same MACF-derived ABD as TipAct and its affinity for actin was measured at about 5 µM by a co-sedimentation assay in MRB40 buffer. Since both ACTIF and TipAct are dimers, this also further validates our assumption that the ACTIF mobile fraction is dimeric. Sequence analysis shows a 73.8% overlap between the ABD of MACF and the ABD of plectin (see sequence alignment in Supplemental Figure 18). Both ABDs consist of two calponin homology (CH) domains in tandem. The isolated CH1 supports binding to actin, albeit with a lower affinity than the tandem; in contrast, CH2 domain has a weaker affinity. The tandem binds at full capacity and has three conserved actin binding sites (Gimona and Winder 1998). The plectin ABD, with its CH domains in a closed conformation, has also been reported to have structural similarity with fimbrin (Sevcik et al. 2004), despite dissimilar sequences. Observations of plectin localization in cells suggest that the high binding affinity for vimentin and low affinity for actin of ACTIF are physiologically relevant. EM images showed that in IF-containing cells, plectin co-localized largely with the vimentin network, whereas in IF-deficient cells, it became mainly associated with the actin cytoskeleton (Foisner et al. 1995).

We showed by electron microscopy that ACTIF induces the formation of crosslinked actin/vimentin bundles and networks already at rather low molar ratios of just 1 ACTIF molecule per 100 actin/vimentin monomers. The isolated IFBD domain could induce bundling of vimentin filaments, but only at much higher (1:1) molar ratios of IFBD:monomer. This finding is consistent with previous work showing that the PRD5 domain by itself can crosslink vimentin (and cytokeratin) filaments (Steinböck et al. 2000). However, we note that contrary to that study, we did not observe any negative effect of the isolated IFBD domain on vimentin filament formation. The fact that the IFBD domain alone can bundle vimentin filaments suggests that it may oligomerize. Indeed in EM images of actin filaments mixed with the IFBD domain, we observed globular structures (non-interacting with F-actin), indicating the presence of IFBD clusters. Oligomerization of full-length plectin has been previously observed by electron microscopy images of rotary-shadowed samples, where filament-bound plectin structures were shown to be oligomeric (Foisner et al. 1988). It would be interesting for future studies to investigate the ultrastructure of the bundles formed by ACTIF crosslinking by thin-section EM or cryo-electron tomography. ACTIF not only forms crosslinks between actin and vimentin filaments, but it also forms crosslinks between actin filaments. This finding is consistent with previous work showing that the ABD of plectin by itself can bundle actin filaments (Fontao et al. 2001). In this earlier study, actin bundling relied on dimerization of the ABD. In the case of ACTIF, we enforced dimerization using a cortexillin coiled-coil spacer domain. We cannot exclude that, additionally, interactions between the ABD domains may also contribute to the observed ACTIF-mediated F-actin bundling. In this study we focused on cytolinker-mediated interactions of vimentin and actin in their filamentous forms. In the future it will be interesting to also test interactions between filamentous and soluble forms (G-actin and vimentin tetramers or unit-length filaments). It was recently shown in cells that the vimentin network obstructs G-actin diffusion, possibly due to transient attractive interactions or binding between the two proteins that contributes to the reduced motion (Wu et al. 2022).

The modular engineered cytolinkers we introduced here can provide a strong basis for understanding the regulation and functions of cytolinker-mediated actin/intermediate filament crosslinking in cells. Using cell-free assays, one can test the interactions of cytolinkers with actin and intermediate filaments under well-controlled conditions. Using cell-based studies, one can then test how these findings carry over to the cellular environment, which is much more complex due to effects such as molecular crowding, nonequilibrium activity, etc. To connect cell-free studies to cell-based studies, it will be important to test the effect of cytolinker size, and more specifically the effect of spacer length. In cell-free studies, the effect of spacer length could be studied by purifying full length plectin or APCn2 or engineered variants with different spacer domains from insect or mammalian cells. In cell-based studies, the effect of spacer length could be studied by transfecting cells with knockouts for endogenous cytolinkers with engineered cytolinker constructs. We note that previous work from our lab with an engineered actin-microtubule crosslinker mimicking ACF7/MACF showed that short constructs with the same spacer domain as ACTIF behaved identically to full-length ACF7/MACF in cell-free assays (Alkemade 2021). Several important open questions could be addressed using our engineered cytolinkers. First, it will be interesting to compare binding and crosslinking by ACTIF and ACTIF-APCn2 with different types of intermediate filaments (IFs), given that the expression pattern of IF proteins is cell-type specific and may convey cell-specific cellular functions. Plectin and other plakins have been shown to possess a broad binding specificity towards different types of IFs, including not only vimentin, which we studied in this work, but also cytokeratins, desmin, GFAP and neurofilaments (De Pascalis et al. 2018, Favre et al. 2011, Foisner et al. 1988, Steinböck et al. 2000), because they recognize the conserved rod domain of IF proteins. By contrast, APC was reported to interact with vimentin and GFAP, but not with keratin, although the structural basis for this is still unknown (Sakamoto et al. 2013). Our system makes it possible to perform quantitative comparisons of the kinetic (un)binding rates and the crosslinking efficiency of different IF types. Second, it is also relevant to uncover how posttranslational modifications (PTMs) of IFs such as phosphorylation (Kraxner et al. 2021), glycosyation (Tarbet et al. 2018) and electrophilic or oxidative modifications (González-Jiménez et al. 2023) may affect interactions with ACTIF and ACTIF-APCn2. Conversely, it would also be interesting to test how PTMs of plectin regulate its interactions with IFs. Cell studies have shown evidence for regulation by phosphorylation of sites like serine 4642 in the C-terminal extremity (Bouameur et al. 2013, Foisner et al. 1996, 1991) and by oxidation and nitrosylation of cysteines in plectin’s IFBD (Spurny et al. 2007). Finally, a recent study showed that the association between plectin and vimentin is mechanosensitive and requires actomyosin contractility (Marks et al. 2022). Our reconstitution assay could be used to test whether this mechanosensitivity is intrinsic to plectin, for instance to its force-sensing plakin domain (Daday et al. 2017).

Our work also provides an interesting proof-of-concept for constructing cytoskeletal systems to be applied in active matter model systems (Berezney et al. 2022, Sanchez et al. 2012) or in synthetic cells (Mulla et al. 2018, Weiss et al. 2018). In these systems, designer crosslinkers like ACTIF could be used to manipulate the material’s mechanical properties and associated dynamics. An interesting expansion of our work would be to engineer a light-activatable cytolinker variant, where the crosslinking effect can be spatiotemporally controlled by illumination. Other interesting modular options to tune cytoskeletal crosstalk are to add tubulin-binding sites or binding sites for actin- or microtubule-based motor proteins.

## 4. CONCLUSIONS

Our results reveal specific crosslinking of actin and vimentin filaments mediated by the engineered crosslinker ACTIF. Actin and vimentin filaments form composite bundles and stiff networks upon crosslinking via ACTIF. This engineered crosslinker enables biochemical, structural, and biophysical studies of crosslinker-mediated cytoskeletal interactions. The crosslinker’s modular design provides an opportunity to test different IF binding domains to better understand intermediate filament binding, as demonstrated by our proof of concept comparing the IFBDs from plectin and APC. This approach also provides the opportunity to unravel the effect that actin-vimentin crosstalk exerts on the mechanical properties of the cytoskeleton, either in a simplified *in vitro* system as reported here, or directly in cells.

## 5. METHODS

### 5.1. Proteins

#### 5.1.1. ACTIF Plasmid Construction

ACTIF was designed to contain an enhanced Green Fluorescent Protein (eGFP) tag followed by the ABD from MACF, the coiled-coiled linker of Cortexillin I, and the IFBD of plectin. We inserted additional residues for flexibility between the different domains, as shown in Supplemental Figure 19. As a vector, we used the TipAct construct (López et al. 2014). For the IFBD insert, we used a plasmid corresponding to residues 4030-4631 aa of the isoform Q9QXS1-1 of mouse plectin, a region that contains the plakin repeat domains 5 and 6 and intervening linker(Nikolic et al. 1996). The two fragments (vector and insert) were amplified using the primers indicated in Supplemental Table S1. PCR amplification was performed using KOD Xtreme Hot start DNA polymerase (#1975). The PCR products were run on a 1% agarose gel (Biorad #1613100EDU) and cleaned using the Wizard^@^ SV Gel and PCR Clean-Up System A9281. For electrophoresis, samples were prepared with TrackIt™ Cyan/Yellow Loading Buffer (Invitrogen #10482035), and TrackIt™ 1 Kb Plus DNA Ladder (Invitrogen #10488085) was used as a ladder. The electrophoresis gel was run at 100V for 20 minutes (Biorad #1613100EDU). Next, the fragments were assembled using the Gibson assembly, following the instructions of NEBuilder HiFi DNA Assembly Cloning Kit (#E5520S). Afterward, the sample was transformed into Dh5α competent *E. coli* cells (#C2987). To determine if the plasmid contained the insert, a miniprep was performed for sequencing via Macrogen. The miniprep was performed using the PureYield™ Plasmid Miniprep System A1222. After sequencing, the plasmid was transformed into *E. coli* BL21 (NEB #C2527).

#### 5.1.2. ACTIF Protein Purification

A pre-culture of *E. coli* BL21 cells expressing ACTIF was grown from a glycerol stock in Luria-Bertani (LB) medium containing 1:1000 diluted kanamycin at 37 °C. Growth was upscaled to a total of 12 liters (separated in 4 cultures of 3 L each), adding 1:1000 diluted kanamycin, and 25 mL of ACTIF preculture. The cultures were incubated at 37 °C in flasks on a shaker platform at 200 rpm. When the Optical Density (OD) reached 0.9, we cooled the 4 cultures down in an ice bath for 30 minutes, then induced overnight expression with 1 mM Isopropyl β-D-1-thiogalactopyranoside (IPTG #11411446001) at 16 °C. The cultures were harvested by a 15 min centrifugation in the Avanti JLA8.1000 fixed angle rotor at 4000 rpm at 25 °C. The supernatant was discarded, and the cell pellets were combined in one 50 mL falcon tube. The cells were resuspended with lysis buffer (20 mM sodium phosphate pH 7.4, 10 % glycerol, 5 mM 2-mercaptoethanol, 500 mM NaCl, 45 mM imidazole), supplemented with cOmpleteTM EDTA-free protease inhibitor cocktail (SigmaAldrich, St. Louis, MO, USA #11836153001) and 1 mg/mL of lysozyme from chicken egg white (Sigma Aldrich #10837059001). The cell solution was incubated on ice for 30 minutes. The lysate was passed through the French press three times at 20 kpsi and next centrifuged at 17, 000 × *g* for 1 hour and 20 minutes at 4 °C (Avanti JXN-26). The supernatant was collected and incubated with 1 mL pre-washed nickel-IMAC resin (Thermo Fisher Scientific #88221) overnight at 4 °C. The beads were pre-washed with wash buffer (20 mM sodium phosphate pH 7.4, 10% glycerol, 5 mM 2-mercaptoethanol, 500 mM NaCl, 45 mM imidazole) in an empty column (Econo-Pac^@^ Chromatography Columns #7321010). The lysate was passed through the disposable column for gravity flow purification. The flow-through was collected and kept at 4 °C. The column was washed three times with wash buffer. To elute the protein from the column, the column was incubated with elution buffer (20 mM sodium phosphate pH 7.4, 10% glycerol, 5 mM 2-mercaptoethanol, 500 mM NaCl, 200 mM imidazole) for 20 minutes at 4 °C. Afterward, an analysis of protein yield and purity for the eluate was performed by SDS-PAGE analysis. To improve the protein purity, a final size exclusion chromatography step was performed on a Superdex 200 10/300 prep grade (Cytiva) column. The column was previously stored in ethanol, and it was washed with 1.5× column volume of MiliQ and then with 1.5× column volume of MRB40 buffer (40*mM* piperazine-N,N*′*-bis(2-ethanesulfonic acid) (PIPES) pH 7.0, 4 mM MgCl2, and 1 mM Ethylene glycol tetraacetic acid (EGTA)). After the column was equilibrated, we injected the sample through the 5 mL loop and ran it at a flow rate of 0.5 mL/min, while collecting 0.3 mL fractions. The fractions of interest as judged from the absorbance at 280 nm were pooled and analyzed by SDS-PAGE gel electrophoresis. The sample was aliquoted at the desired molar concentration, calculated using values for the molecular weight and extinction coefficient of 138 kDa and 101,245 M^−1^ cm^−1^, respectively, which were theoretically estimated with Expasy analysis tool (Walker 2005). The aliquots were stored at −80 °C. We typically obtained 7 mg from 12 L of bacterial cell culture at a concentration of about 7 mg/mL.

By quantifying the intensity of the protein bands observed on the SDS-page gel in Supplemental Figure 3 using the plugin called “Gels” on Fiji, we estimated the purity of ACTIF at 64%. To test whether the lower-molecular weight bands are degradation products of ACTIF, we performed fluorescence imaging of the SDS-page gels using a Typhoon (Supplemental Figure 4). The most prominent bands in the SDS-page gel displayed a fluorescent GFP signal, indicating that these bands indeed result from partial proteolysis of ACTIF. Western blot analysis where we stained the bands with anti-GFP antibodies confirmed this conclusion (Supplemental Figure 5). Finally, we also performed liquid chromatography–mass spectrometry (LC–MS) to assess the purify of the ACTIF preparation. LC-MS analysis showed that the most abundant protein in the sample had a molecular weight of 138 kDa, as expected from the amino-acid sequence of ACTIF (Supplemental Figure 6). Putative identification of the main peaks of the LS-MS chromatogram allowed us to identify degraded fragments of ACTIF sample.

#### 5.1.3. IFBD(plectin) Construction

The IFBD(plectin) construct consisted of an N-terminal His6-tag followed by the region of the isoform Q9QXS1-1 of mouse plectin of 4030-4631 aa, corresponding to the plakin repeat domains 5 and 6 and intervening linkers, as shown in Supplemental Figure 20. The plasmid assembly was performed as reported for ACTIF, and the used primers are presented in Supplemental Table S3.

#### 5.1.4. ACTIF-APCn2 Construction

ACTIF-APCn2 was designed as a variant of ACTIF, with the plectin-derived IFBD replaced by the armadillo repeats of APC, again using the TipAct construct (Alkemade et al. 2022, López et al. 2014) as a vector. For the insert, we used a plasmid courtesy of Sandrine Etienne-Manneville that corresponded to the APCn2 fragment of APC, specifically from residue 334 to 740 (Sakamoto et al. 2013). The assembly was performed as reported for ACTIF using the primers presented in Supplemental Table S2. The final sequence is shown in Supplemental Figure 21.

#### 5.1.5. IFBD(plectin) protein purification

IFBD(plectin) was purified following the same protocol as for ACTIF, but no gel filtration was required. In order to calculate the final concentration of the purified sample, the respective molecular weight and extinction coefficients are 71 kDa and 48,165 M^−1^ cm^−1^, which were calculated based on the protein sequence with the Expasy analysis tool (Walker 2005). We typically obtained 5 mg from 6 L of bacterial cell culture.

#### 5.1.6. ACTIF-APCn2 protein purification

ACTIF-APCn2 was purified following the exact same protocol as ACTIF until the lysis step. Culture pellets were resuspended in 40 mL of lysis buffer (20 mM sodium phosphate pH 7.4, 500 mM NaCl, 45 mM imidazole, 5 mM bME, 10 % v/v glycerol), 1 cOmplete™, EDTA-free Protease Inhibitor Cocktail (Roche) and 1 mg/mL of lysozyme from chicken egg white (Sigma Aldrich). Lysis of the spun down cultures was performed by sonication, using a 422A tip, with a protocol following amplitude 40 %, 10 seconds on, 10 seconds off, for a duration of 10 minutes. Afterwards, the lysate was centrifuged using TI45 40,000 rpm, for 30 min at 4 °C. The supernatant was loaded on a 5mL HisTrap FF column (Cytiva) and the purification was performed using the AKTA system. The column was washed with 50 mL lysis buffer and was eluted with 50 mL lysis buffer supplemented with 200mM imidazole. The flow rate of the buffers was 1 mL/min. Fractions with the clean protein based on the SDS-gel were pooled. In order to calculate the final concentration of the purified sample, the respective molecular weight and extinction coefficients are 116 kDa and 94,770 M^−1^ cm^−1^, which were theoretically estimated with Expasy analysis tool (Walker 2005). We typically obtained 6-8 mg from 12 L of bacterial cell culture at a concentration of 6-8 mg/mL. By quantifying the intensity of the protein bands observed on the SDS-page gel, we estimated a purity of 72% (Supplemental Figure 3).

#### 5.1.7. Actin purification

For EM and rheology experiments, lyophilized rabbit alpha-skeletal muscle actin was purchased from Hypermol EK. Actin was dialyzed against G-buffer (pH 7.8) comprising 5 mM Tris-HCl, 0.1 mM MgCl_2_, 0.2 mM ATP and 5 mM DTT. After dialysis, G-actin was aliquoted, flash-frozen in liquid nitrogen and stored at −80 °C. The final actin concentration was measured by UV-VIS absorbance measurements using an extinction coefficient at 290 nm of 26,600 M^−1^ cm^−1^ and molecular weight of G-actin of 42 kDa (Houk Jr and Ue 1974).

For TIRF experiments, alpha-skeletal actin was purified from rabbit muscle acetone powder following a protocol in ref. (Wioland et al. 2017) that was based on an earlier protocol (Spudich and Watt 1971). G-actin was fluorescently labeled on accessible surface lysines using ATTO-643 succinimidyl ester (Life Technologies) as described previously (Romet-Lemonne et al. 2018). Similarly, actin was labeled with biotin using biotin succinimidyl ester (Life Technologies). An SDS-page gel showing the purity of the actin preparation is provided in Supplemental Figure 3.

#### 5.1.8. Vimentin purification

Wild-type human vimentin was purified from TG1 *E. coli* bacteria (Sigma-Aldrich) following an earlier protocol (Herrmann et al. 2004). In brief, we started the induction when the OD reached 1.2, and cultured the bacteria in Terrific Broth medium overnight at 37 °C. Then, we collected the bacterial cells by centrifuging the culture medium, and lysed them with lysozyme, together with DNase (Roche), RNase (Roche), and protease inhibitors (Pefabloc and PMSF) in a 50 mM TRIS buffer pH 8. We washed the released inclusion bodies 5 times in the presence of DTT, and protease inhibitors: first with 200mM NaCl, 1% sodium desoxycholate, 1% NP40, 20mM TRIS pH 7.5, 2mM EDTA; second with 10mM TRIS pH 8, 0.5% Triton X 100; third with 10 mM TRIS pH 8, 0.5% Triton X 100 and 1.5M KCL; fourth with 10 mM TRIS pH 8, 0.5% Triton X 100; and finally with 10 mM TRIS, 0.1 mM EDTA pH 8 and 20µl DTT. Washing includes successive steps of incubation on ice for 20 min, centrifugation at 8000 rpm for 10 min at 4 degrees Celsius, and resuspension using a cooled douncer. After washing, we resuspended the inclusion bodies in a denaturing buffer (8 M urea, 5 mM Tris pH 7.5, 1 mM EDTA, 1 mM DTT, 1 % PMSF) and centrifuged them at high speed (100000 × *g*) for 1 h. Upon collecting the supernatant, we conducted vimentin purification by two sequential steps of exchange chromatography, using first an anionic (DEAE Sepharose, GE Healthcare) and then a cationic (CM Sepharose, GE Healthcare) column in a buffer of pH 7.5 containing 8 M urea, 5 mM TRIS, 1 mM EDTA, and 0.1 mM EGTA. We collected the vimentin protein in 2 mL Eppendorf tubes and monitored the protein concentration by Bradford assay. Only the most concentrated fractions were selected and pooled together. We stored the vimentin at −80 °C with an additional 10 mM methylamine hydrochloride solution. An SDS-page gel showing the purity of the vimentin preparation is provided in Supplemental Figure 3.

For experiments, about 100 µL denatured vimentin aliquots were transferred to dialysis tubing (Servapor, molecular weight cut off at 12 kDa) and vimentin was renatured by stepwise dialysis from 8 M, 6 M, 4 M, 2 M, 1 M, 0 M urea into sodium phosphate buffer (pH 7.0, 2.5 mM sodium phosphate, 1 mM DTT) at room temperature with at least 15 minutes for each step. The final dialysis step was performed overnight at 4 °C in 2 L of the sodium phosphate buffer. The vimentin concentration was determined by UV-VIS absorbance measurements using a molecular weight of 53.65 kDa and extinction coefficient at 280 nm of 22,450 M^−1^ cm^−1^ (Aufderhorst-Roberts and Koenderink 2019).

#### 5.1.9. Fluorescent labeling of vimentin

We labeled vimentin proteins by coupling an Alexa Fluor 555 dye to cysteine-328, as described previously (Tran et al. 2023, Winheim et al. 2011). In short, denatured vimentin stored in 8 M urea was dialyzed for 3 h in a labeling buffer of pH 7.0 containing 50 mM sodium phosphate and 5 M urea. Next, the fluorescent dye (AF-555 C2 maleimide, ThermoFisher) dissolved in DMSO was added in a molar ratio of 5:1 of (dye:vimentin). The solution was gently mixed for 1 h at room temperature, and the reaction was then quenched by the addition of 1 mM DTT. We used a Dye Removal Column (#22858, ThermoFisher) to remove excess dye. Finally, we renatured the labeled vimentin by stepwise dialysis from 8 M, 6 M, 4 M, 2 M, 1 M, 0 M urea to sodium phosphate buffer (pH 7.0, 2.5 mM sodium phosphate, 1 mM DTT). The final product was stored at 4 °C and used within 10 days.

#### 5.1.10. SDS-page gels

We used 10% Mini-PROTEAN TGX precast polyacrylamide gels with 10 wells (Biorad). We loaded 10 µL of proteins at 10 µM mixed in a 3:1 ratio with 4X Laemmli Buffer Sample (Biorad) provided with 10% DTT. We ran the gels for 1h at 120 V with Tris/Glycine/SDS buffer (Biorad) and stained the gels with InstantBlue protein stain (Expedeon) for 1h. To asses purity before and after gel filtration of ACTIF, we used a 4-15% Mini-PROTEAN TGX Precast Protein Gels, 15-well, 15uL (BIORAD). We loaded 10 µL of protein mixed in a 1:1 ratio with Laemmli 2x Concentrate (Merck). The gel was ran for 35min at 200V with Tris/Glycine/SDS buffer (BIORAD) and stained with InstantBlue Coomassie Protein Stain (Cat # Ab119211) for 2 hours. The intensity of the bands observed on the SDS-page gels were analyzed using the plugin “Gels” from Fiji (Schindelin et al. 2012).

#### 5.1.11. Typhoon imaging

An SDS-PAGE gel with the samples of interest was prepared. The proteins were diluted with 2x Laemmli Sample buffer (2x Laemmli Sample Buffer S3401-1VL). 10 µL of proteins (at concentrations of 53 µM for ACTIF, 30 µM for GFP and 122 µM for VCA) were loaded onto the Mini-Protean TGX Gels. The gels were ran for 35 min at 200 V in 1x Tris/Glycine/SDS buffer from Bio-Rad. Gels were afterwards washed with demi-water and imaged with the Amersham Typhoon. The wavelengths for Cy2, Cy3 and Cy5 were chosen, according to the marker (Colorcoded Prestained Protein Marker Broad Range (10-250kDa) *Lot*4 74154S)). Images of the three wavelengths were combined using ImageQuant TL.

#### 5.1.12. Western blot

The samples were boiled at 95 °C for 5 min in 3:1 Laemmli sample/loading buffer provided with 10 % DTT. These samples were spun down for 5 min at 16000× *g* before loading 10 µL into 10 % Mini-PROTEAN TGX precast gels (Bio-Rad). The gel was run on gel electrophoresis at 150 V for 1 h. After the run was complete, we performed a wet transfer to a nitrocellulose membrane (Bio-Rad) for 1 h at 0.09 A at room temperature. The membrane was then blocked with 4% milk solution diluted in TBS 1X provided with 0.5 % Tween 20 (TBS-T) for 30 min. After the blocking, the membrane was incubated with 10 mL of anti-GFP primary antibodies (Roche, Cat.#11814460001) at a concentration of 1 µg/mL in 4 % milk/TBS-T for 1 h. The membrane was then washed three times for 15 min with TBS-T on a shaker. For the secondary antibody staining, the membrane was incubated with 10 mL TBS-T with secondary antibodies Mouse HRP at a dilution of 1:5000 (Jackson ImmunoResearch, Cat.#715-035-151) for 45 min at room temperature on a shaker. After incubation, the membrane was washed 3 times for 15 min in TBS-T. We used a ChemiDoc Imaging System (Bio-rad) to image the western blot. Activation of the HRP was done using an ECL Western Blotting Substrate detection kit (Pierce, Cat.#PI32106).

#### 5.1.13. Liquid chromatography–mass spectrometry (LC–MS)

Intact protein analysis by LC-MS was performed as described previously Lignieres et al. (2023). Samples were injected onto a custom made BioResolve RP (450 Å, 2.7 µm, 0.3 mm × 150 mm) Polyphenyl Column (Waters, Milford, MA, USA) using an Acquity M-Class chromatographic system from Waters. Mobile phases A and B were H_2_O and H_2_O/acetonitrile 20/80, respectively, acidified with 0.1 % (v/v) Difluoroacetic Acid. Samples were eluted at a flow rate of 6 µL/min using the following slope change points: 0-5 min hold at 12.5 % B, 10 min gradient to 25 % B, 40 min gradient to 50% B, 10 min wash at 80 % B and finally 5 min hold at 12.5 % B. The eluent was sprayed using the conventional ESI ion source of a Waters Cyclic IMS SELECT SERIES mass spectrometer operating in the positive ion mode. 500ng were injected on the column. The time of flight was operated in V-mode with a resolution specification of 60,000 FWHM at a mass range of m/z 50-4000.

#### 5.2. Total Internal Reflection Fluorescence Microscopy (TIRF-M)

#### 5.2.1. Assembly of fluorescent vimentin and actin filaments

Prior to the experiment, we polymerized vimentin filaments from AF-555-labeled vimentin (with 10 % labeling fraction) at a concentration of 3.7 µM in V-buffer (2.5 mM sodium phosphate, pH 7.0, 100 mM KCl) for 3 h at 37 °C using a water bath. We also polymerized F-actin from 10 µM G-actin labeled with ATTO-643 (10 % labeling fraction) in F-buffer (20 mM Tris–HCl, pH 7.4, 50 mM KCl, 2 mM MgCl_2_, 0.5 mM ATP) for 1 h at room temperature. In experiments that required F-actin immobilization, we assembled biotinylated F-actin from 10 µM G-actin labeled with both ATTO-643 (10 % labeling fraction) and biotin (1 % labeling fraction) in F-buffer for 1 h at room temperature. We note that labeling fractions of 10 % do not impact the assembly of actin (Jégou et al. 2011) nor vimentin (Tran et al. 2023).

#### 5.2.2. TIRF-M experimental setup

We assembled the composites of vimentin, F-actin and ACTIF in a flow chamber. The flow chamber was constructed from silanized coverslips following a previously described protocol (Leduc et al. 2012, Tran et al. 2023). Briefly, we incubated cleaned coverslips with dichlorodimethylsilane 0.05 % in trichloroethylene for 1 h at room temperature. We sonicated the coverslips 3 times in methanol, each for 15 min, to wash off the excess silane. We then heated up two silanized coverslips compressing pieces of parafilm in between to 50 °C on a hot plate. This made the parafilm melt and attach to the two coverslips, resulting in a flow chamber with a volume of about 10 microliters.

For the control experiment to test the actin-vimentin direct interaction, we first flowed vimentin antibodies (SC666260, Santa Cruz Biotechnology) at a concentration of 2 µg/mL in F-buffer into the chamber and incubated for 5 min. We passivated the flow chamber by flowing Pluronic F127 1 % in F-buffer and incubating for 15 min. We continued flowing bovine serum albumin (BSA) 5 % in F-buffer and incubated for 15 min. Finally, we thoroughly rinsed the chamber by flowing an extensive amount of F-buffer. We prepared ATTO-643-labeled G-actin (10 % labeling fraction) at 1 µM in F-buffer supplemented with an oxygen scavenger mixture (glucose 1.2 mg/mL, glucose oxidase 40 µg/mL, glucose catalase 8 µg/mL, DTT 1 mM) and 0.2 % methylcellulose 4000 cP, and gently flowed them into the chamber with vimentin filaments on substrate. We captured images of the whole process of F-actin polymerizing from G-actin and rapidly elongating to long filaments while freely moving close to the vimentin filaments.

For the TIRF-M assay in which vimentin filaments were attached to the substrate, we used a similar protocol to have the flow chamber coated with vimentin antibodies and passivated with F127 and BSA as described in the control experiment above. We then began the composite assembly by diluting pre-assembled vimentin filaments 100 times in F-buffer (final concentration: 37 nM), then flowing them into the chamber and incubating for 5 min for filament attachment onto the vimentin antibodies. Next, the chamber was rinsed, and ACTIF diluted in F-buffer supplemented with the oxygen scavenger mixture to a final concentration of 5.3 nM was flushed into the chamber while imaging had started in advance in order to capture the early state of protein attachment to vimentin filaments. After 15 min, we performed FRAP assay by photo-bleaching a region of ACTIF binding to vimentin filaments and observing the fluorescence recovery over 20 min. Finally, pre-assembled actin filaments were diluted 100 times in F-buffer with an oxygen scavenger (final concentration: 0.1 µM) and flushed into the chamber, forming vimentin-ACTIF-actin composites.

For the assay with immobilized F-actin, we flowed neutravidin (Cat. #31000, ThermoFisher) at a concentration of 10 µg/mL in F-buffer into the chamber and incubated for 5 min. We again passivated the flow chamber with Pluronic F127 1 % in F-buffer and BSA 5 % in F-buffer, and thoroughly rinsed the chamber afterward, similar to the previous assay. Here, our first step in the composite assembly was diluting pre-assembled biotinylated F-actin 100 times in F-buffer (final concentration: 0.1 µM), then flushing them into the chamber, and incubating for 5 min. F-actin with biotin was constrained on the substrate due to the neutravidin coating. Then, we rinsed the chamber and flowed in diluted ACTIF at 5.3 nM or at 530 nM in F-buffer with oxygen scavenger while imaging the binding reaction for 15 min. We also performed a FRAP assay by photo-bleaching a region of ACTIF on F-actin and recording the recovery for 20 min. Finally, pre-assembled vimentin filaments diluted 100 times in F-buffer with oxygen scavenger (final concentration: 37 nM) were flowed into the chamber for the formation of actin/vimentin composite bundles.

TIRF microscopy was performed on a Nikon Eclipse T*i* inverted microscope controlled by MicroManager, equipped with a 60× oil-immersion objective and a Kinetix sCMOS Camera (Photometrics). The TIRF illumination was performed using an Ilas2 module (GATACA Systems). The experiment temperature was maintained at 25 °C using objective and microscope stage temperature controllers.

### 5.3. Interferometric Scattering Microscopy

Interferometric Scattering Microscopy (iSCAT) experiments were conducted with a OneMP Mass Photometer (Refeyn). The instrument was equipped with a 525 nm laser for illumination. iSCAT measurements were carried out using CultureWell gaskets (Cat. #GBL103250, Grace Biolabs) affixed to #1.5 coverslips (Cat. #13296788, Corning). To prepare the coverslips for optimal protein adhesion, a meticulous stepwise sonication cleaning process was employed. The coverslips were sonicated sequentially in MilliQ water, 50% isopropanol, and again in MilliQ water, each for a duration of 5 minutes. Following the cleaning process, the coverslips were incubated with a poly-L-lysine solution (PLL, Cat. #P4832, Merck) for 30 seconds. Subsequently, the coverslips were rinsed with MilliQ water and dried under a stream of N_2_ gas. The treated coverslips were stored upright in a Teflon rack within a covered beaker to protect them from dust. They were utilized within a week to maintain their cleanliness. During the iSCAT measurements, videos were recorded with a field of view measuring 10 × 10 µm. The recording duration ranged from 6000 to 15000 frames, with a frame rate of 300 frames per second. This ensured a minimum count of 1000 particles per video. Maintaining a dilute solution of proteins is crucial to ensure well-separated scattering patterns with no overlap. Hence protein concentrations (in the range of 10 nM to 50 nM, dilution with F-buffer from the stock concentration of the protein) were chosen to achieve adequate separation of landing events while still collecting sufficient statistics within a maximum of 15000 frames to limit data volume. The iSCAT video analysis was executed using the DiscoverMP software, a commercial software package provided by Refeyn.

### 5.4. Transmission Electron Microscopy

Electron microscopy analysis was conducted on samples deposited on Cu400 carbon support grids (Quantifoil, Cat. #N1-C73nCu40-01) with a JEM-1400Plus transmission electron microscope (JEOL) operating at an acceleration voltage of 120 kV. Imaging was carried out using a TemCam-F416 CMOS camera (TVIPS) with a resolution of 4k×4k pixels. The sample preparation protocol was as follows: Initially, the Cu400 grids underwent a glow discharge treatment in an oxygen plasma using the GloQube-D instrument provided by Quorum Technologies Ltd. This procedure was essential to enhance protein adsorption. Subsequently, a 4 µL droplet of the protein sample was delicately deposited onto the grid and allowed to adsorb for 1 to 2 minutes. To eliminate any surplus protein and salt, the samples underwent three washed with MilliQ water with careful blot-drying after each wash. Finally, the samples were stained for 25 seconds with a 2% aqueous solution of uranyl acetate from Electron Microscopy Sciences. Following staining, the samples were meticulously blot-dried once again to eliminate any remaining liquid. We made two independent repeats per sample condition. For each independently prepared sample we usually prepared 2 grids (in case one would be broken or overstained). To study the samples of vimentin and IFBD, we took a total of 100 images depicting all conditions. To study the samples of actin, vimentin and ACTIF, we took a total of 180 images depicting all conditions. To study the samples of actin, vimentin and ACTIF-APCn2, we took a total of 100 images depicting all conditions. Many bundles were imaged twice, since we looked at different magnifications to inspect different regions of the bundles in detail. Upon measuring bundle widths for statistics, we carefully avoided measuring the same bundle.

### 5.5. Rheology

Rheological measurements were conducted using a stress-controlled KinexusMalvern Pro rheometer equipped with a stainless steel cone-plate geometry. The cone had a radius of 20 mm and an angle of 1°. The temperature was maintained at 37 °C by Peltier plates. F-actin-vimentin networks were polymerized between the plates of the rheometer by loading 40 µl of the sample directly after mixing the proteins into the polymerization buffer (F-buffer). To prevent solvent evaporation during the experiment, a thin layer of mineral oil Type A was carefully applied around the sample edge. Progress of network formation was monitored by applying a small oscillatory shear with a fixed strain amplitude of 0.5% and an oscillation frequency of 0.5 Hz for 2 hours. Following the 2-hour polymerization period, a frequency sweep was conducted in the range of 0.01 to 10 Hz at a small strain amplitude of 1%, sampling 10 data points per decade. Data taken at frequencies exceeding 3 Hz had to be discarded because they were dominated by inertial effects from the rheometer. For each curve, we took an average over *N* = 2 independently prepared samples.

### 5.6. Data analysis

All quantitative analysis of TIRF imaging data was performed using Fiji. The images were pre-processed with background subtraction and image registration. To quantify colocalization of F-actin, vimentin, and the cytolinkers, we used the coloc2 plugin from Fiji to perform a pixel intensity correlation over space. To retrieve reaction rates of ACTIF on vimentin filaments (or on F-actin), we manually extracted the intensity values over time in the ACTIF channel along 30 to 50 vimentin filaments. In this experiment, we performed 3 independent repeats with one field of view for each repeat. In each repeat, we calculated the mean and standard deviation of ACTIF intensity over 30 - 50 ROIs and fitted the intensity curves using GraphPad Prism. To retrieve the number of actin filaments attached to vimentin through ACTIF, we quantified the pixel fluorescence intensities in the F-actin channel along vimentin filaments (~150 filaments, *N* = 2 independent experiments) and plotted the distribution of intensities. This distribution displayed multiple peaks, with the first peak at an intensity of ~600 a.u., as expected for single F-actin in these imaging conditions. The distribution was fitted by a multiple Gaussian, with the peak positions constrained to be multiples of the position of the first peak, and common width for all the peaks except the last one.

To obtain the unbinding rate of ACTIF from vimentin filaments (or from F-actin) from FRAP data, we again first performed background subtraction and image registration. Then, we again selected 30 ROIs following shapes of vimentin filaments (or F-actin) from a pre-bleached image. The selected filaments needed to be inside the photo-bleaching region. Then we extracted the intensity value of ACTIF over time from the selected ROIs. In this experiment, we also performed 3 independent repeats with one field of view for each repeat. In each repeat, we obtained the mean and standard deviation of the ACTIF intensity over 30 ROIs and fitted the intensity curves using GraphPad Prism.

For TEM microscopy, image analysis was performed by Fiji. In order to determine the filament widths, intensity profiles perpendicular to the segment of interest were plotted at multiple locations along the filament contours. The filament diameter was determined from the width of the bright area between the darker borders. For copolymerized vimentin and IFBD, *N* = 28 bundles were measured for the 1:10 IFBD:vimentin molar ratio and *N* = 17 bundles for the 1:1 IFBD:vimentin molar ratio. For copolymerized actin and vimentin with ACTIF, *N* = 40 bundles were measured for each ACTIF:monomer ratio.

For shear rheology, data was extracted from the rSpace Software (NETZSCH Analyzing and Testing). Custom Python scripts were used to plot *G′* and tan(δ) as a function of oscillation frequency. We calculated an average of *N* = 2 curves of independently prepared experiments per condition.

## CREDIT AUTHORSHIP CONTRIBUTION STATEMENT

IIP: Conceptualization, Methodology, Validation, Investigation, Formal analysis, Visualization, Writing – original draft preparation; QDT: Conceptualization, Methodology, Validation, Investigation, Formal analysis, Visualization, Writing – original draft preparation; AG: Methodology, Validation, Investigation; ZM: Methodology, Validation, Investigation; GW: Conceptualization, Writing – review and editing; CL: Conceptualization, Methodology, Validation, Investigation, Formal analysis, Writing – original draft preparation; Supervision, Project administration, Funding acquisition; GHK: Conceptualization, Methodology, Validation, Investigation, Formal analysis, Writing – original draft preparation; Supervision, Project administration, Funding acquisition.

## DATA AVAILABILITY

Data will be made available on request.

## ACKNOWLEDGEMENTS

The authors would like to thank Jeffrey de Haan for protein purification and Eli van der Sluis for great teaching and discussions about cloning techniques. We also would like to thank Sonam Marapin for the Western blot analysis, as well as Guillaume Chevreux, Jean-Michel Camadro and Marie Ley, from the IJM proteomics platform, for the LS-MS analysis. We thank Sandrine Etienne-Manneville for the APCn2 construct and the entire Romet-Lemonne/Jégou lab for actin purification and labeling as well as technical help and stimulating discussions. We gratefully acknowledge funding from the NWO Talent Programme which is financed by the Dutch Research Council (project number VI.C.182.004), the Centre National de la Recherche Scientifique, and the Agence Nationale de la recherche (project number ANR-21-CE11-0004). QDT was supported by the LabEx “Who am I?” (ANR-11-LABX-0071) and the Université Paris Cité IdEx (ANR-18-IDEX-0001), funded by the French Government through its “Investments for the Future” program.

## SUPPLEMENTARY MATERIAL

### SUPPLEMENTARY TABLES AND FIGURES

**Supplemental Table 1:**
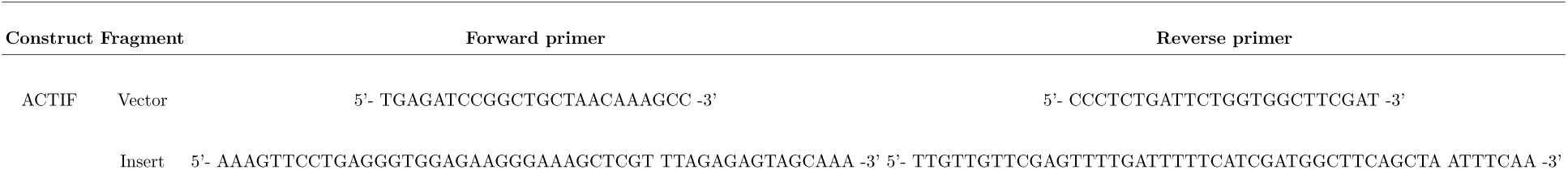
Primers used for Gibson Assembly of ACTIF construct.

**Supplemental Table 2:**
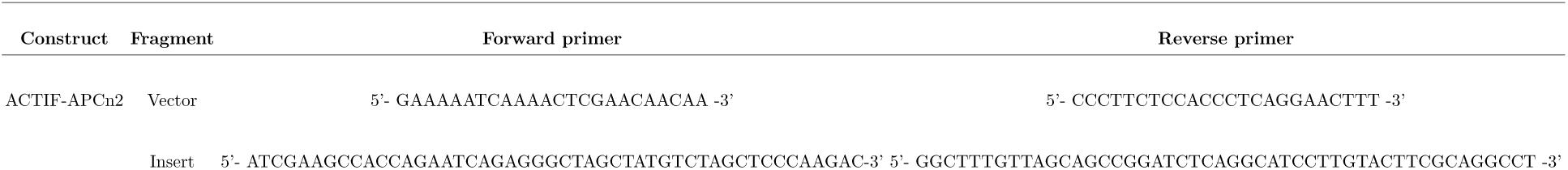
Primers used for Gibson Assembly of ACTIF-APCn2 construct.

**Supplemental Table 3:**
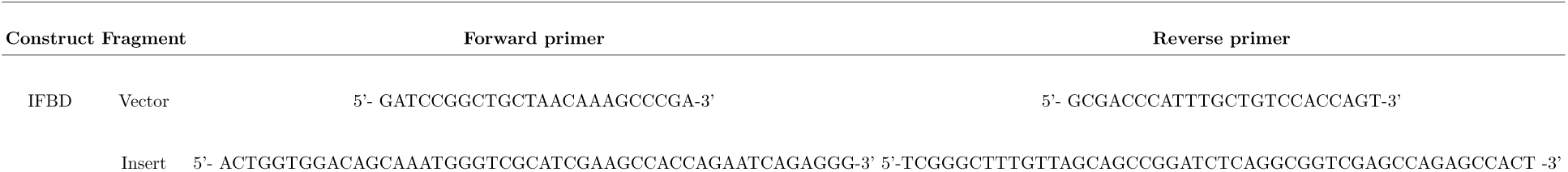
Primers used for Gibson Assembly of IFBD(plectin) construct.

**Supplemental Figure 1:**
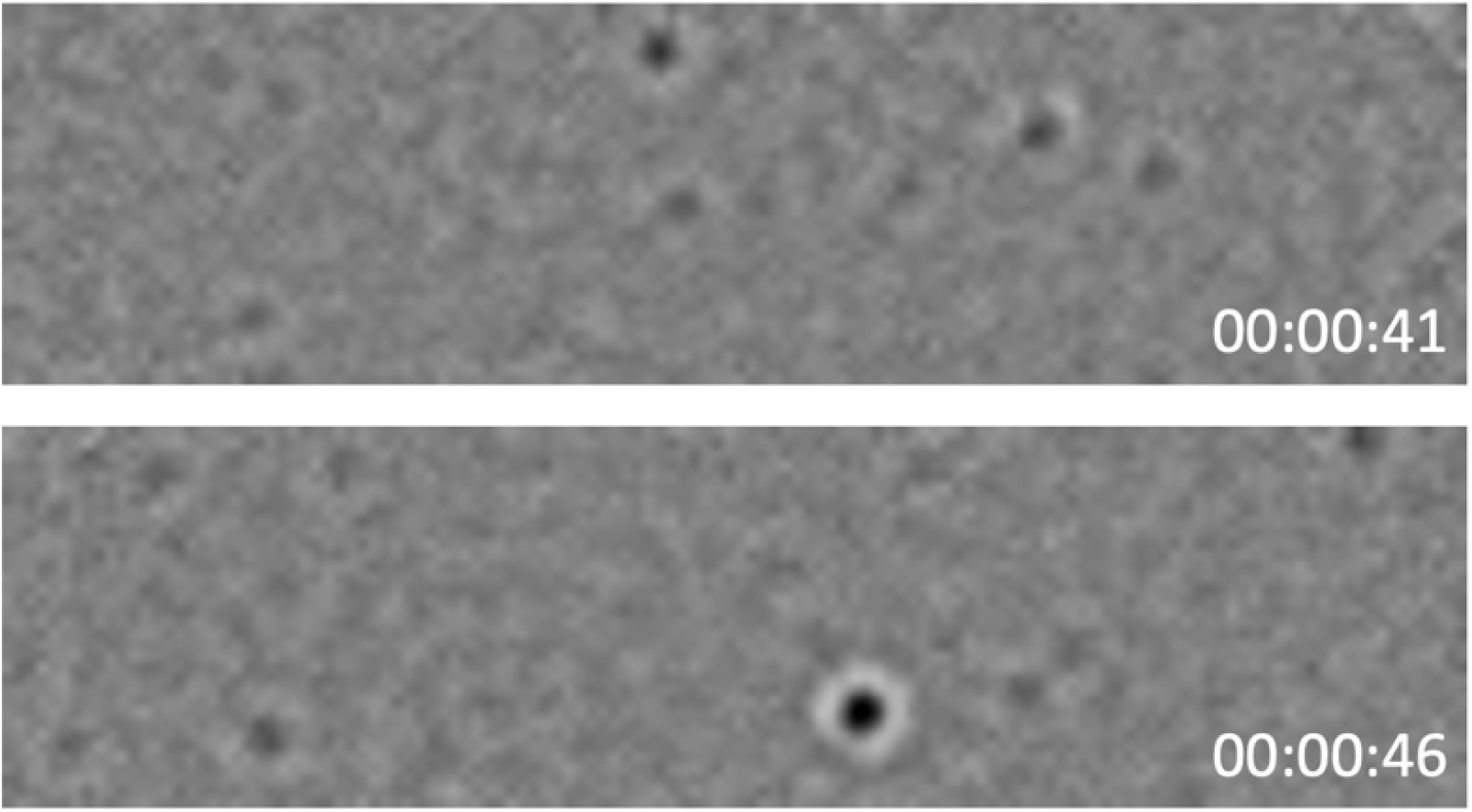
Frames from a time-lapse video recorded for interferometric scattering microscopy of a sample of 40nM ACTIF. Different time frames are shown in seconds. The area of detection shown is 18 µm^2^

**Supplemental Figure 2:**
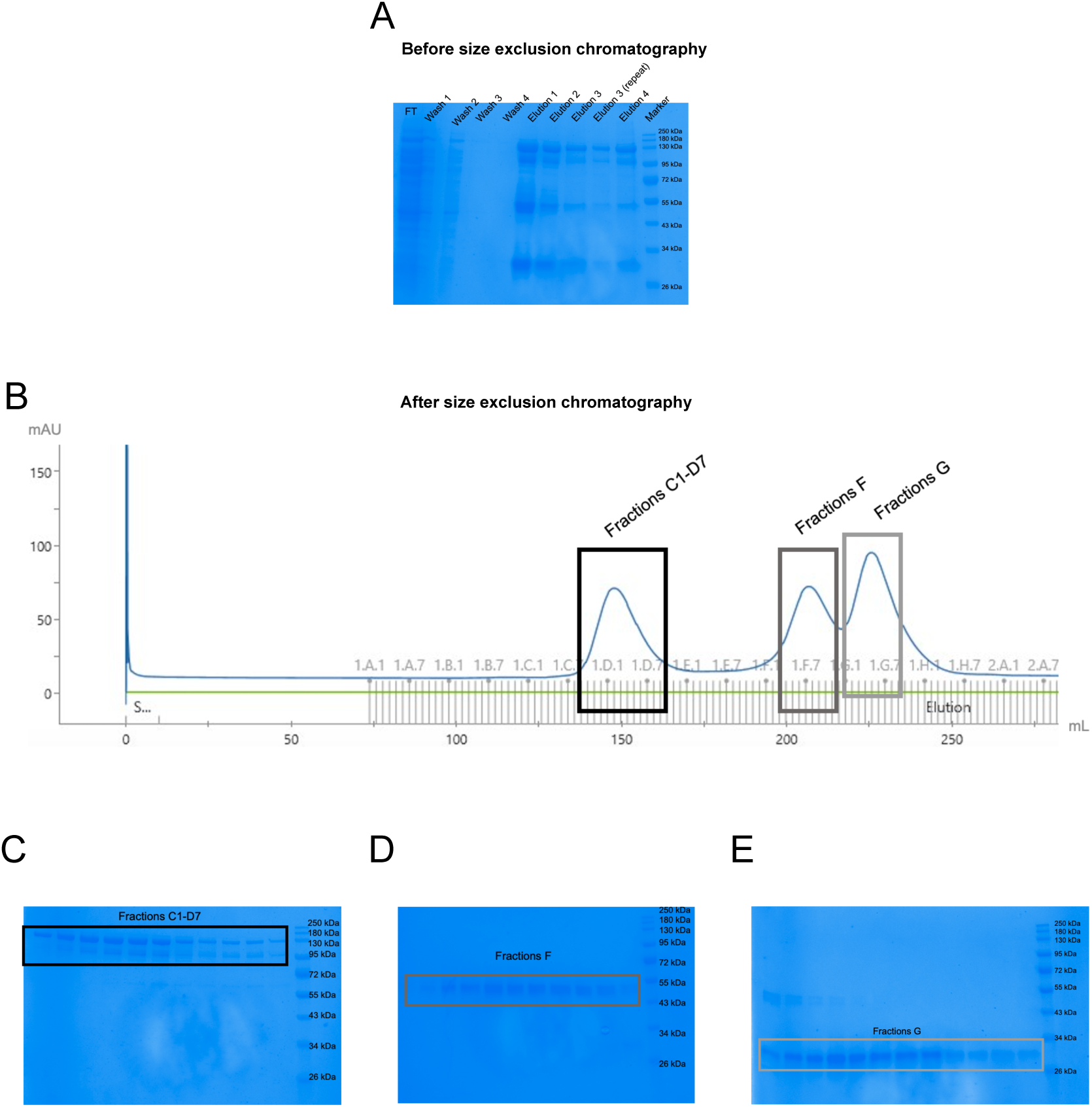
Gel filtration purification of ACTIF. (A) SDS-PAGE gel of ACTIF fractions obtained by His-tag protein purification. The lanes show the flowthrough (FT), fractions from the wash steps, and fractions from the elution steps. On the right, the molecular weight reference. (B) Chromatogram of the size exclusion purification with the pooled fractions (C1-D7 fractions, F fractions and G fractions) marked by the rectangles. The first peak corresponds to the expected molecular weight of ACTIF (138 kDa). (C) SDS-PAGE gel of the pooled fractions C1-D7, corresponding to the first peak in the chromatogram. (D) SDS-PAGE gel of the pooled fractions F, corresponding to the second peak in the chromatogram. (E) SDS-PAGE gel of the pooled fractions G, corresponding to the third peak in the chromatogram.

**Supplemental Figure 3:**
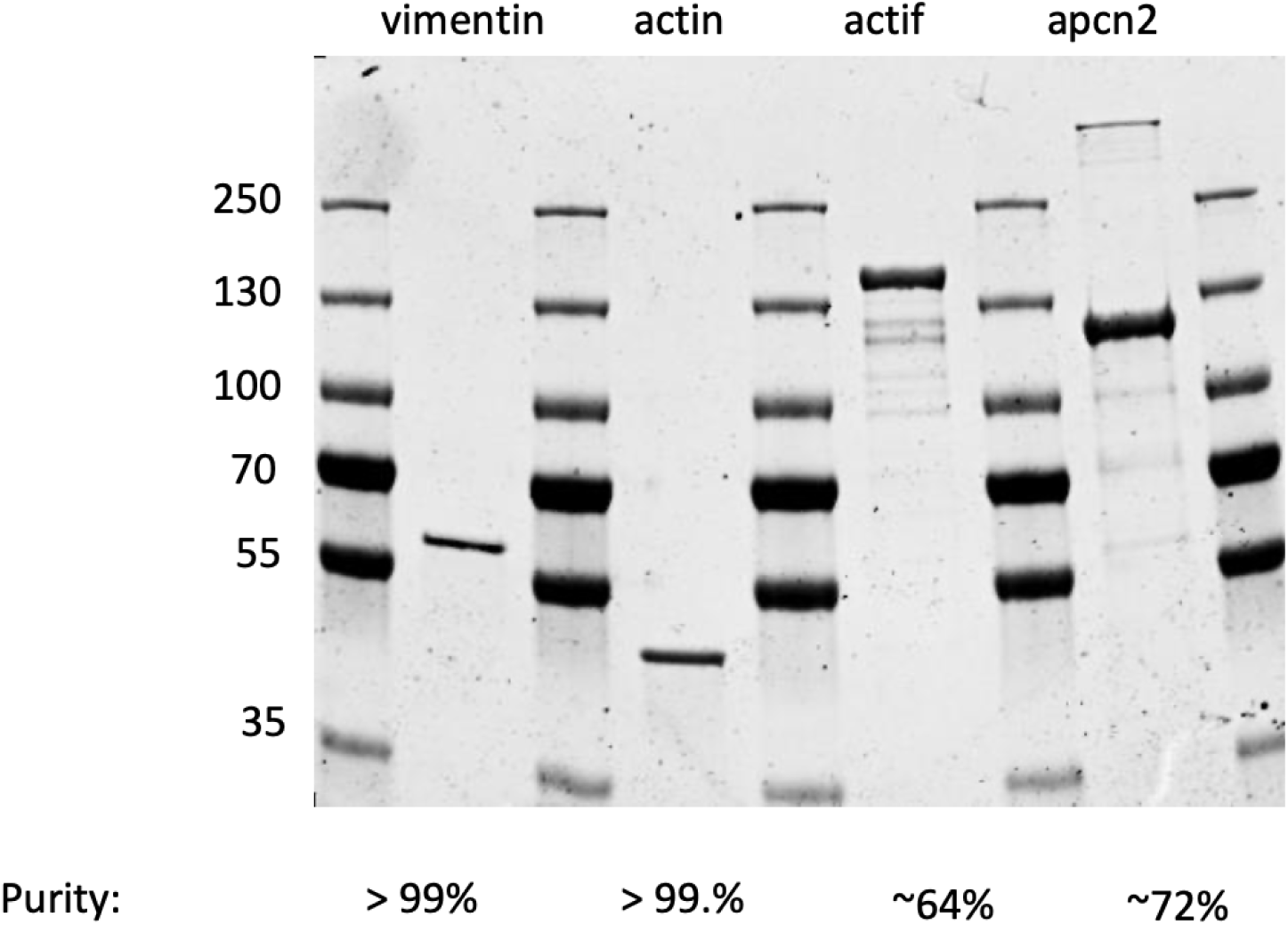
SDS-PAGE gel of purified vimentin, G-actin, ACTIF and ACTIF-APCn2. The purities of the full-length products as determined by densitometry analysis of the gel are specified at the bottom.

**Supplemental Figure 4:**
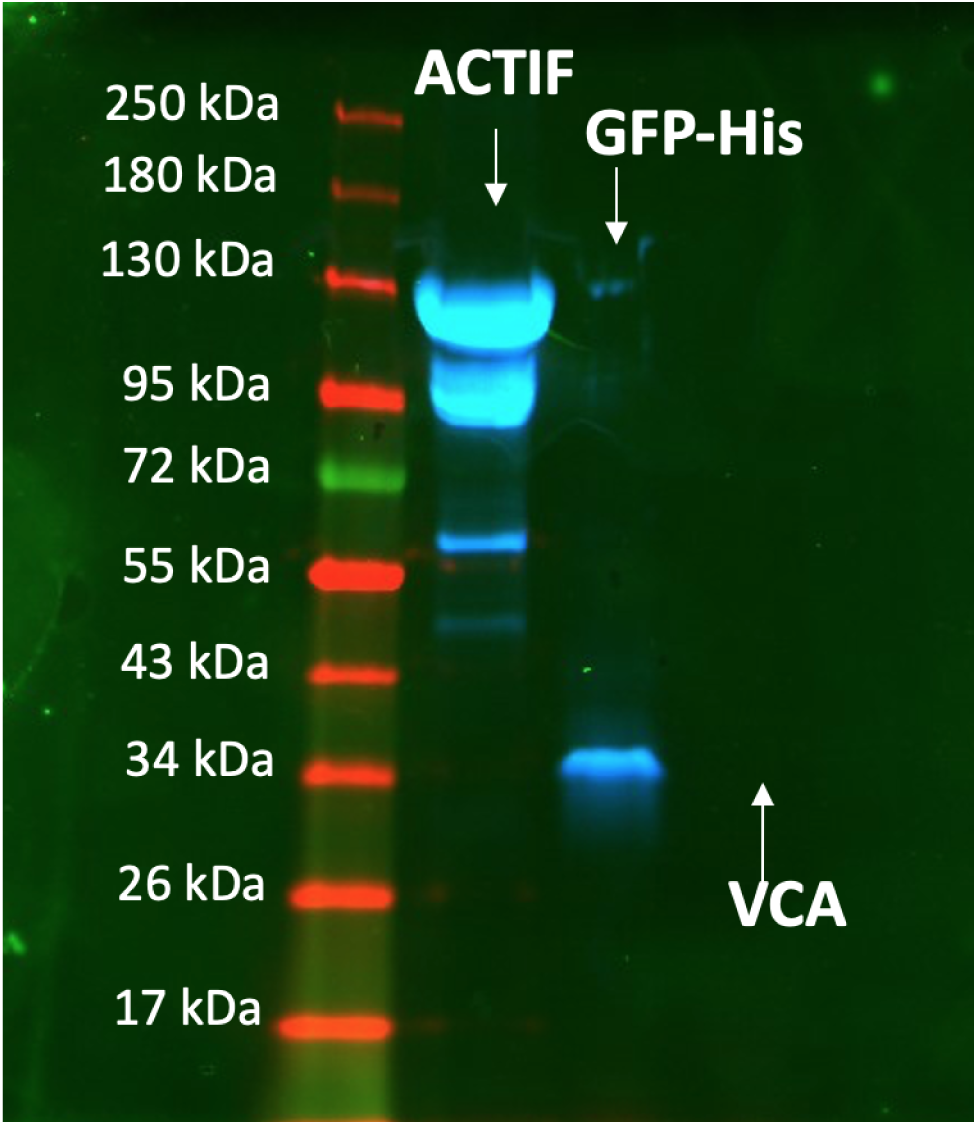
Typhoon fluorescence imaging of an SDS-PAGE gel containing ACTIF, His-tagged GFP (as a positive control) and VCA (non-fluorescent control protein). Green fluorescent signal is detected for full length ACTIF and the lower bands (indicating proteolysis) and for the His-tagged GFP control sample. No fluorescent signal is detected for VCA.

**Supplemental Figure 5:**
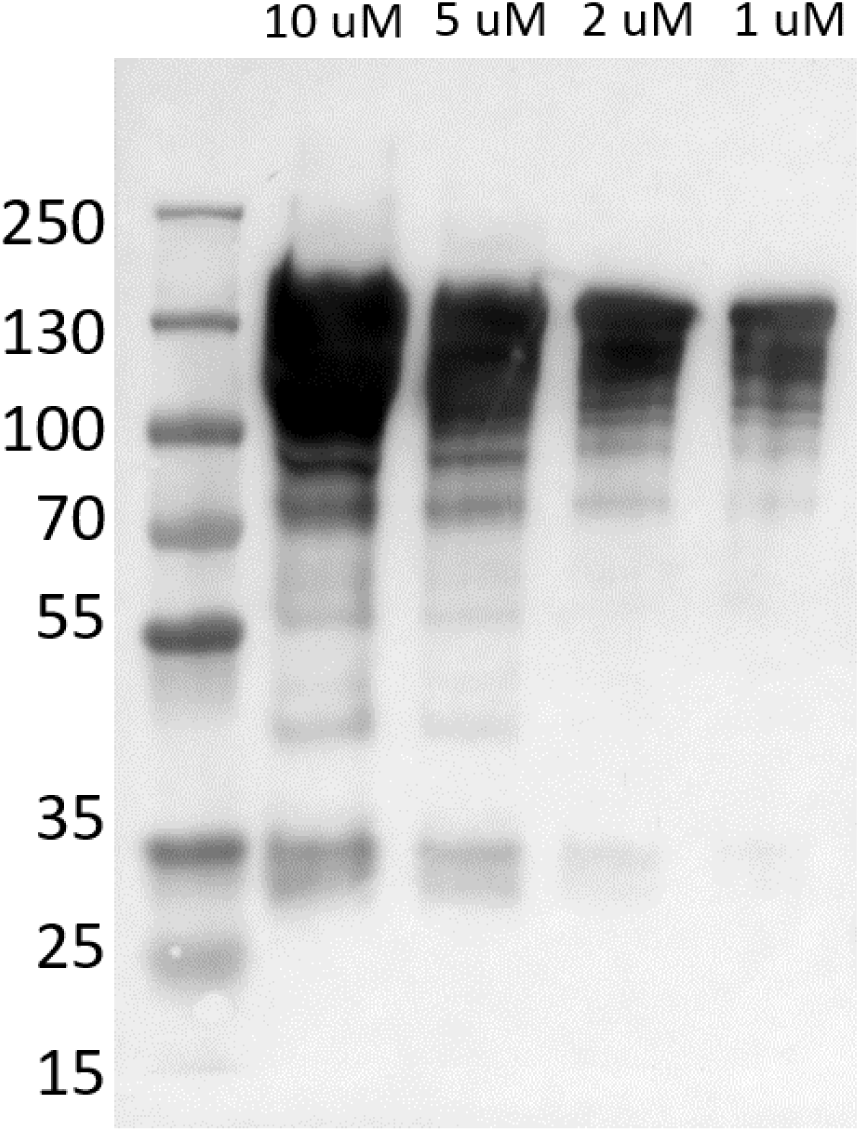
Western blot analysis of an ACTIF preparation with mouse anti-GFP antibodies. Blot imaging of activated secondary antibody labeled with HRP overlayed with a colorimetric image of the protein ladder. The different lanes correspond to different ACTIF dilutions, and the protein concentration is written on top of the lanes. 10 µL of protein/Laemli was loaded by lane.

**Supplemental Figure 6:**
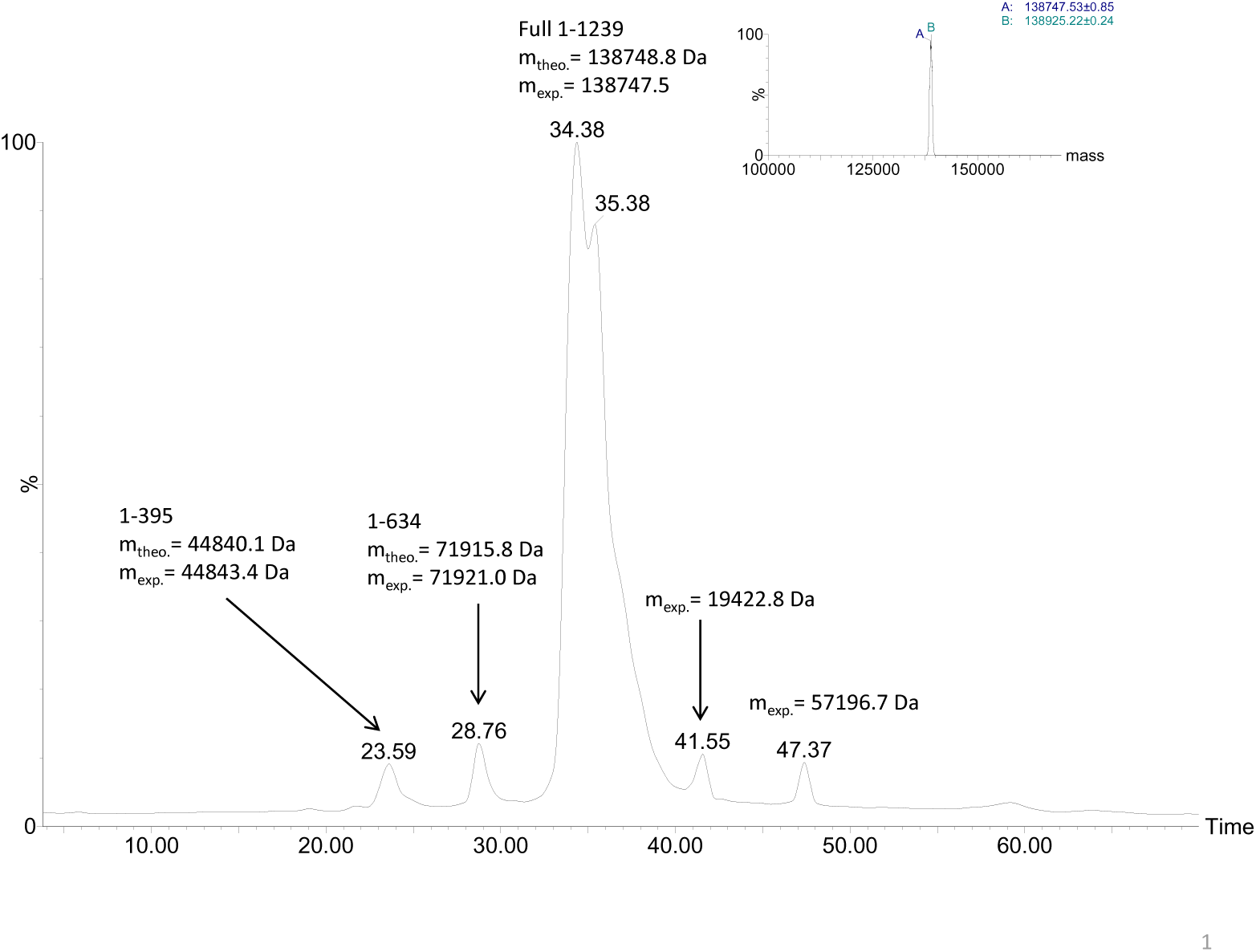
Liquid chromatography–mass spectrometry (LC–MS) analysis of ACTIF. Putative identification of the peaks in the mass spectrogram is indicated on the graph with the size of the corresponding ACTIF fragment, the theoretical mass and experimental mass. Inset: Deconvolved mass spectra of the most abundant proteins with masses of 138747.5 Da and 138925.2 Da, corresponding to full length ACTIF with N-gluconoylation (+178 Da), a standard post-translational modification obtained for N-terminally His-tagged proteins expressed in E.Coli.

**Supplemental Figure 7:**
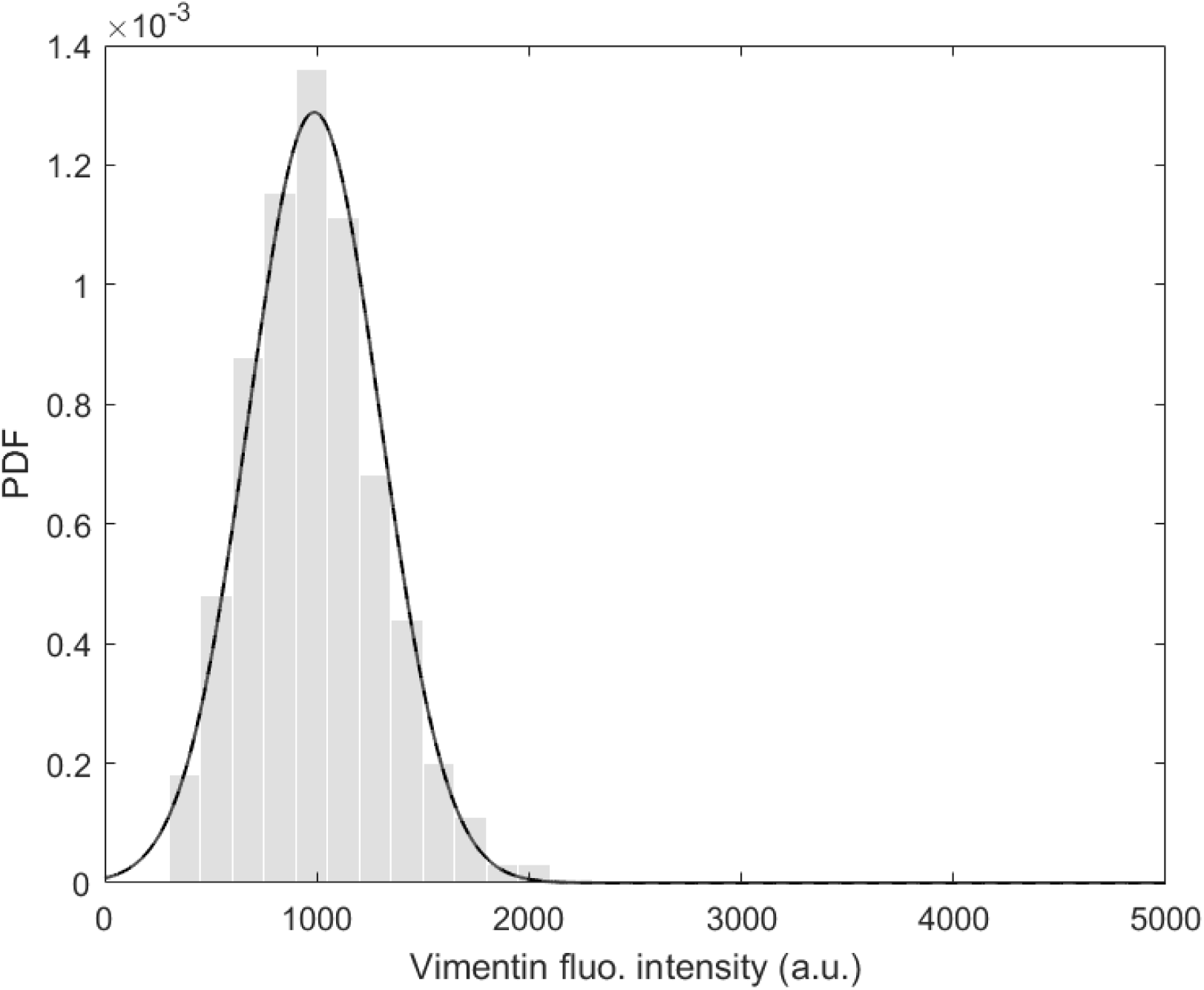
Distribution of pixel fluorescence intensities along vimentin filaments. The distribution appears as a single peak with the average intensity expected for single filaments under these imaging conditions, indicating that vimentin exists mainly as single filaments when attached to the substrate. Sample size, 100 filaments. PDF, Probability density function.

**Supplemental Figure 8:**
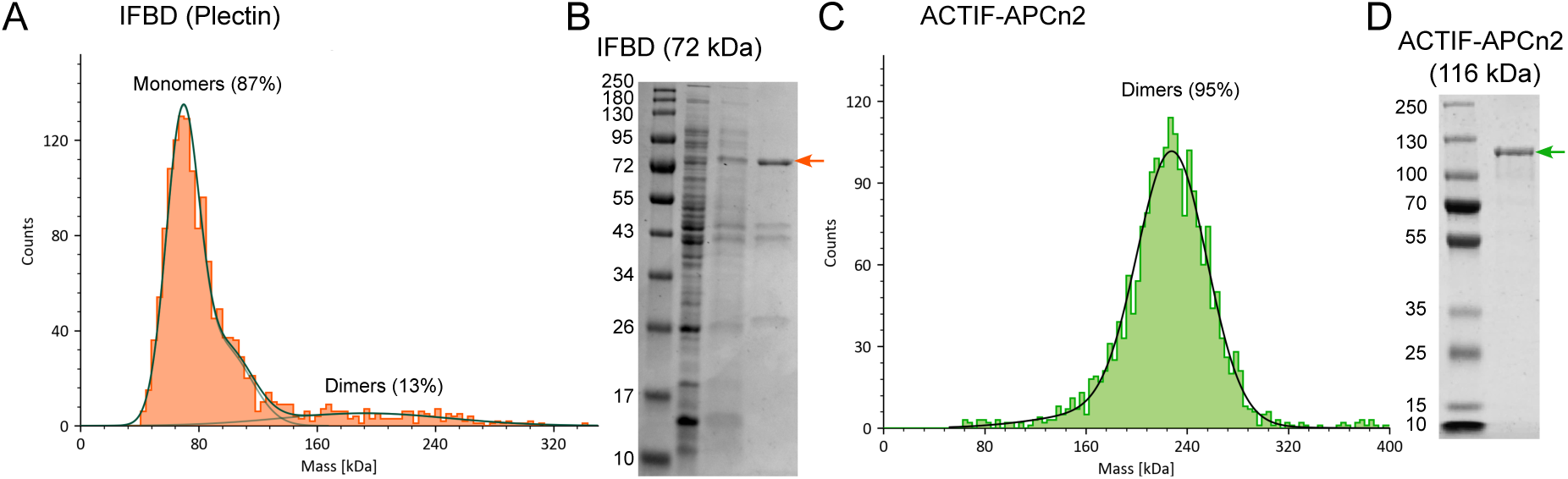
iSCAT and SDS-PAGE of ACTIF-APCn2 and IFBD. (A) iSCAT histogram count of IFBD (plectin), portraying a mostly monomeric population with a 13% dimeric population count. (B) SDS-PAGE of the pooled fractions of IFBD(plectin) after purification, with an arrow marking the band of interest. (C) iSCAT histogram count of APCn2, portraying a 95% dimeric population. (D) SDS-PAGE of the pooled fractions of APCn2, with an arrow marking the band of interest. For the gels, 10 µL of proteins at 10 µM were loaded and stained with Coomassie.

**Supplemental Figure 9:**
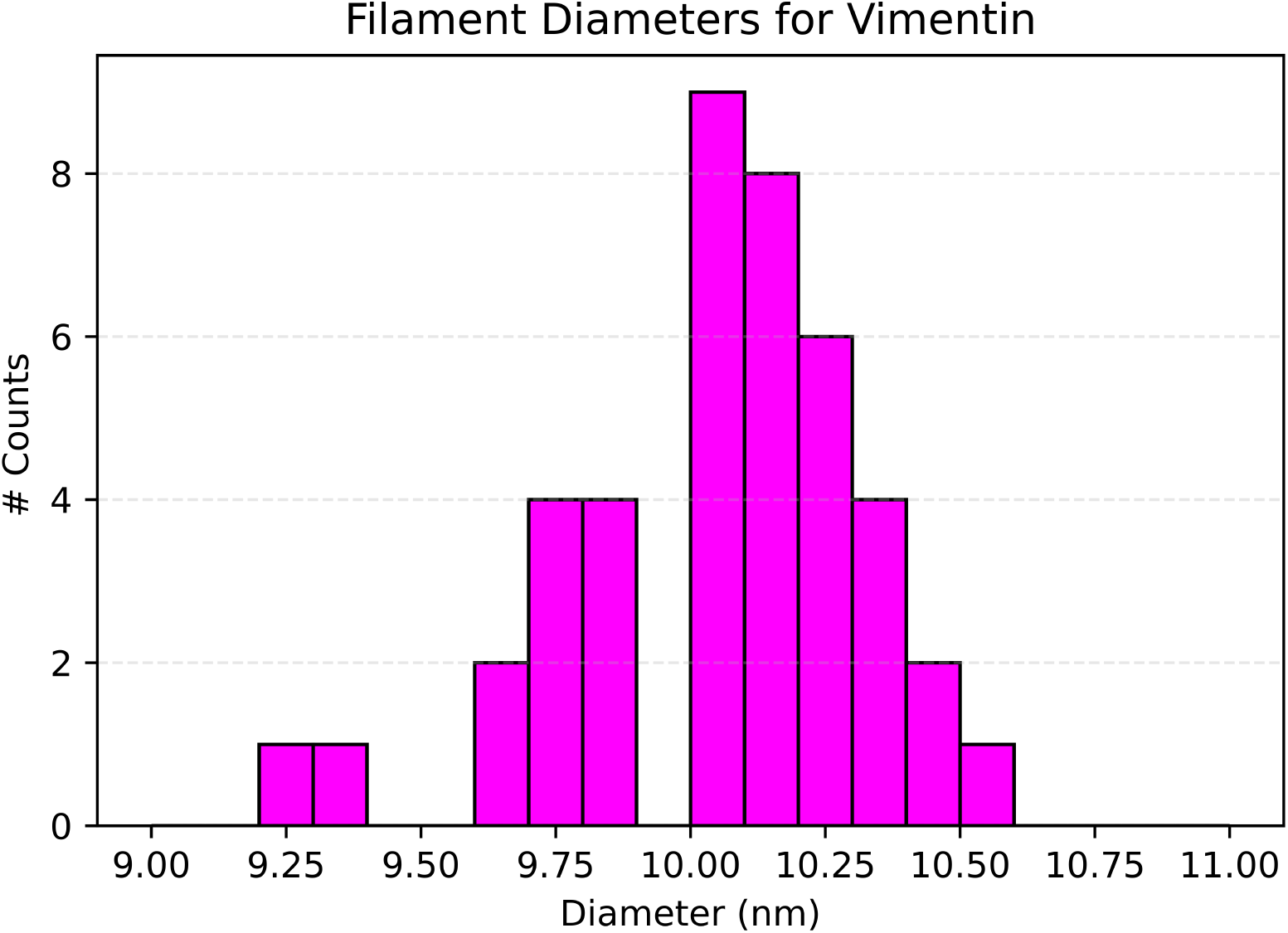
Histogram of diameters of randomly sampled vimentin filaments (*N* = 50).

**Supplemental Figure 10:**
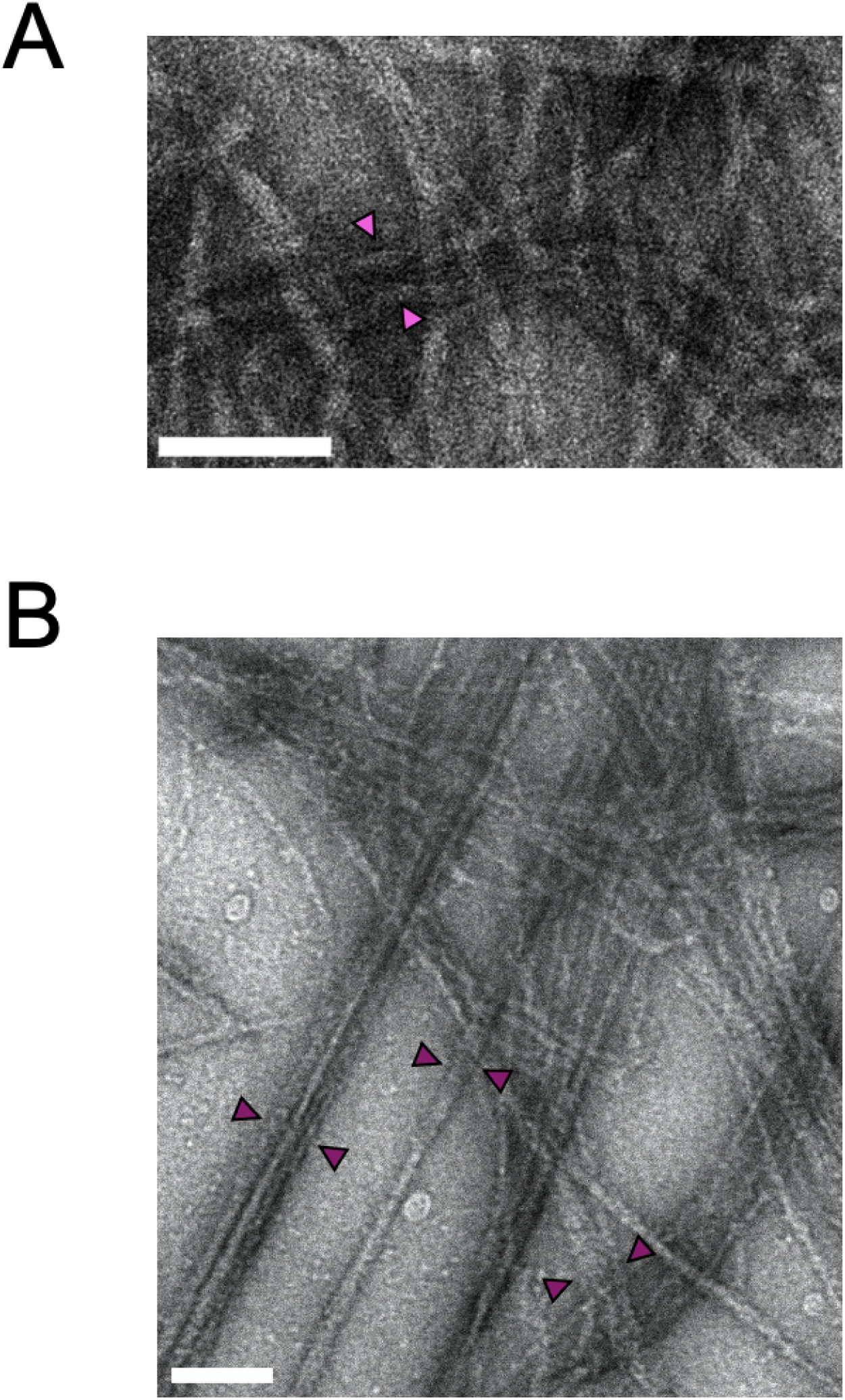
Zoomed-in transmission electron microscopy images of 4 µM of vimentin filaments bundled with (A) 0.4 µM IFBD or (B) 4 µM IFBD. The arrows indicate examples of measured bundles. Scale bars indicate 100 nm.

**Supplemental Figure 11:**
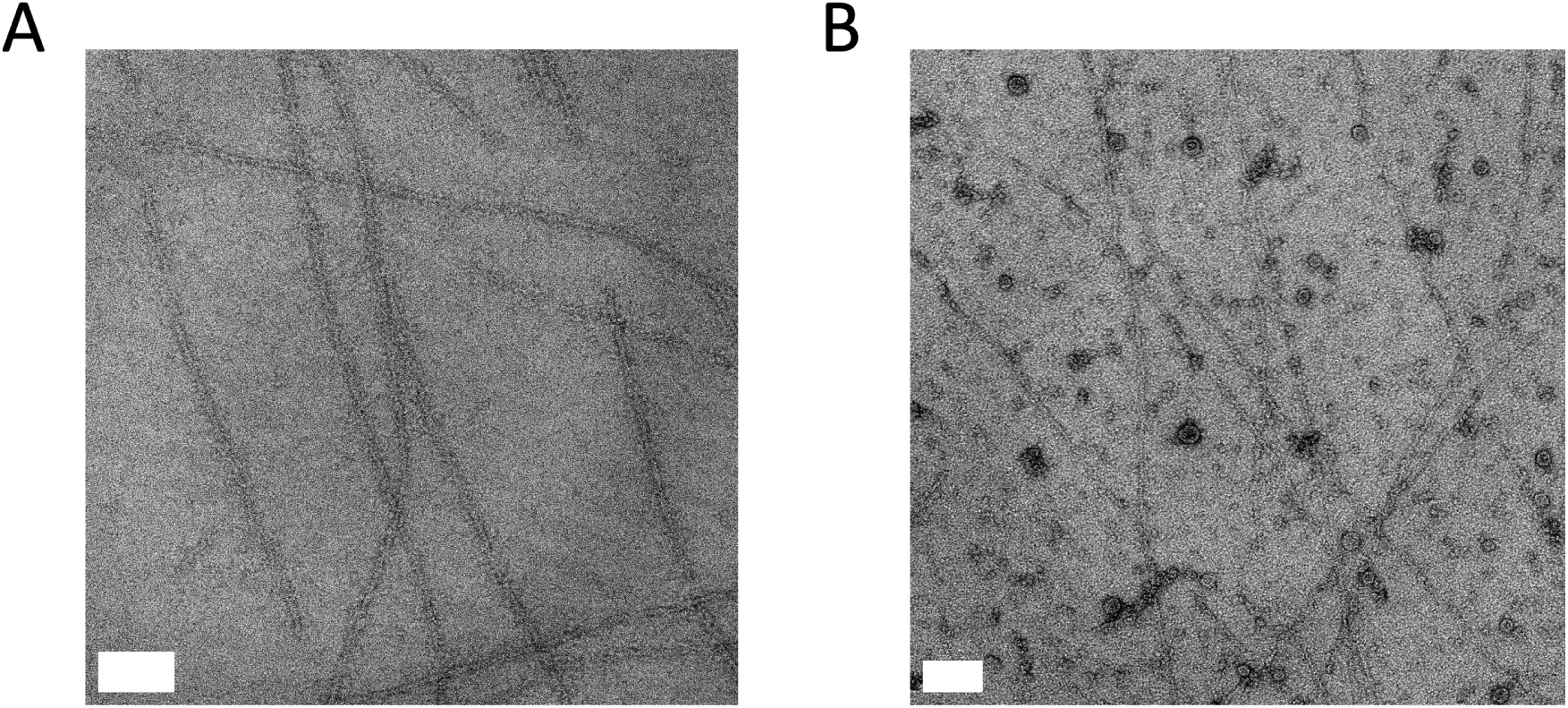
Transmission electron microscopy images of actin filaments at 2 µM concentration (A) polymerized without IFBD or (B) with 20 *mu*M IFBD (from plectin). The IFBD tends to oligomerize (see globular structures present in (B) but not in (A)). We do not observe evidence for any interactions between actin and the globular IFBD structures. Scale bars indicate 100 nm.

**Supplemental Figure 12:**
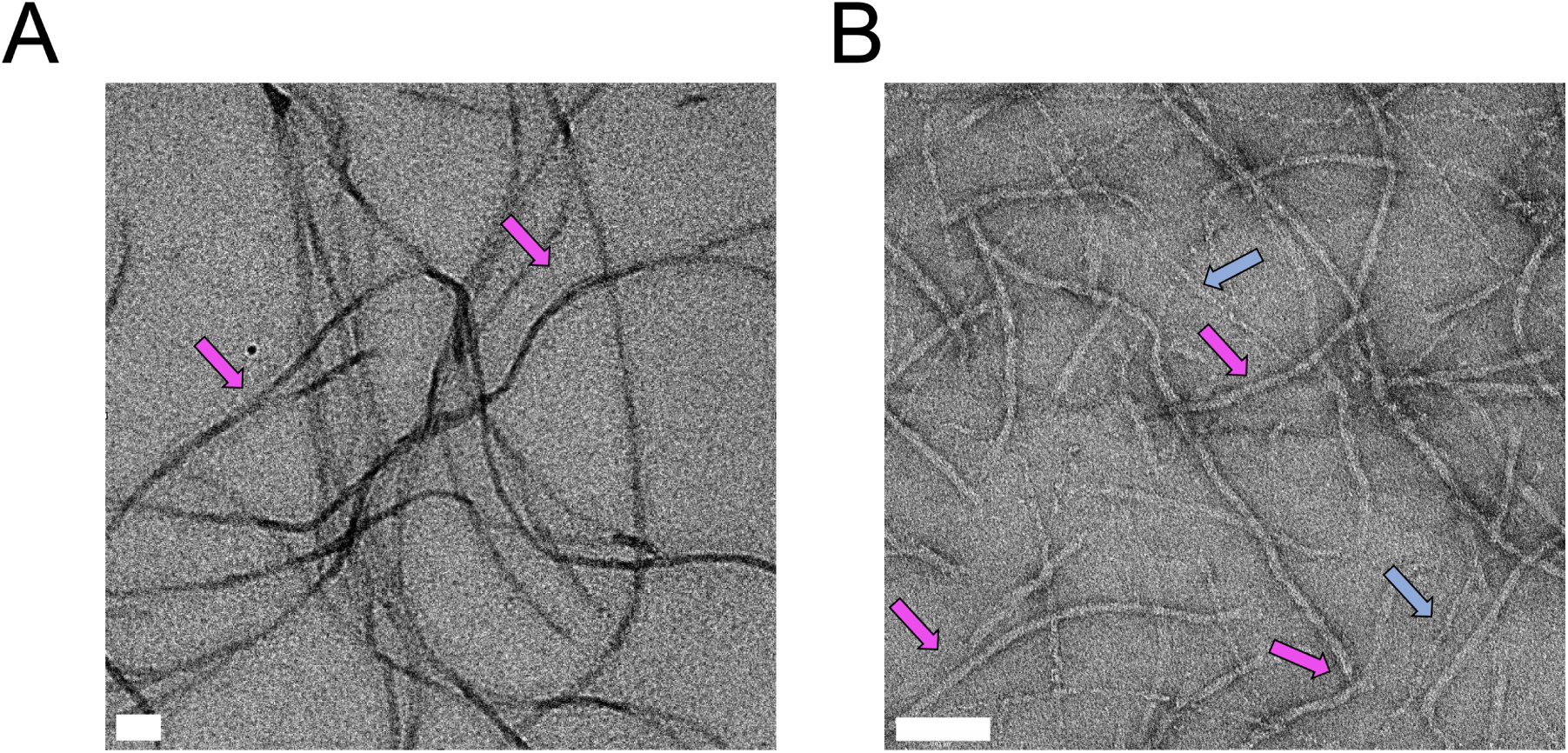
Zoomed-in electron microscopy images comparing vimentin and co-polymerized vimentin-actin composites. (A) 4 µM vimentin filaments. (B) 1 µM vimentin and 1 µM F-actin. Pink arrows indicate vimentin filaments, which are recognizable based on their larger diameter and smaller persistence length as compared to F-actin. Blue arrows in B indicate F-actin. Scale bars indicate 100 nm.

**Supplemental Figure 13:**
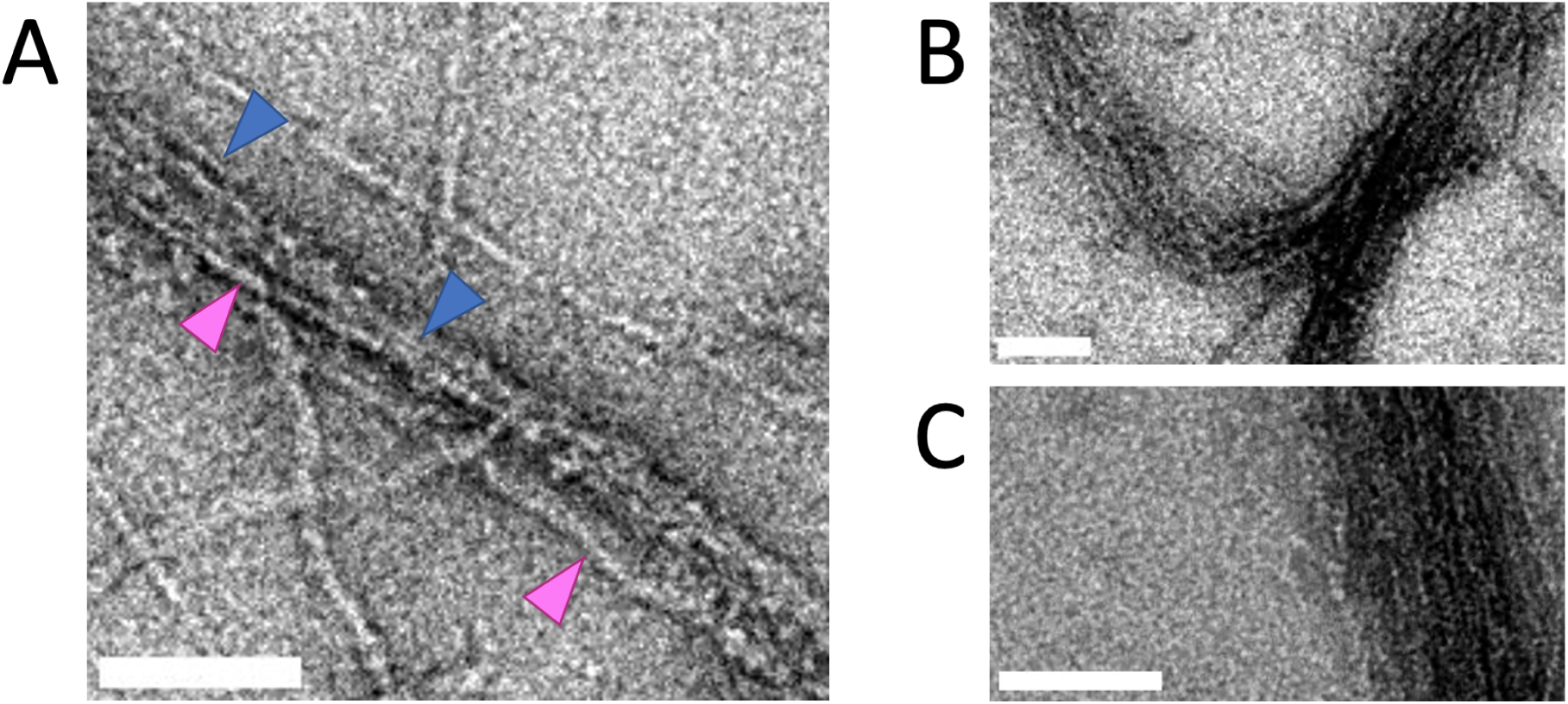
Transmission electron microscopy of copolymerized actin, vimentin and ACTIF showing detailed views of the bundles. (A) Mixed actin and vimentin within a bundle (blue arrows for actin, magenta for vimentin). (B) Another example, showing variations in bundle packing along the bundle length. (C) Bundle where we only observe actin filaments, based on their diameter. Scale bars indicate 100 nm.

**Supplemental Figure 14:**
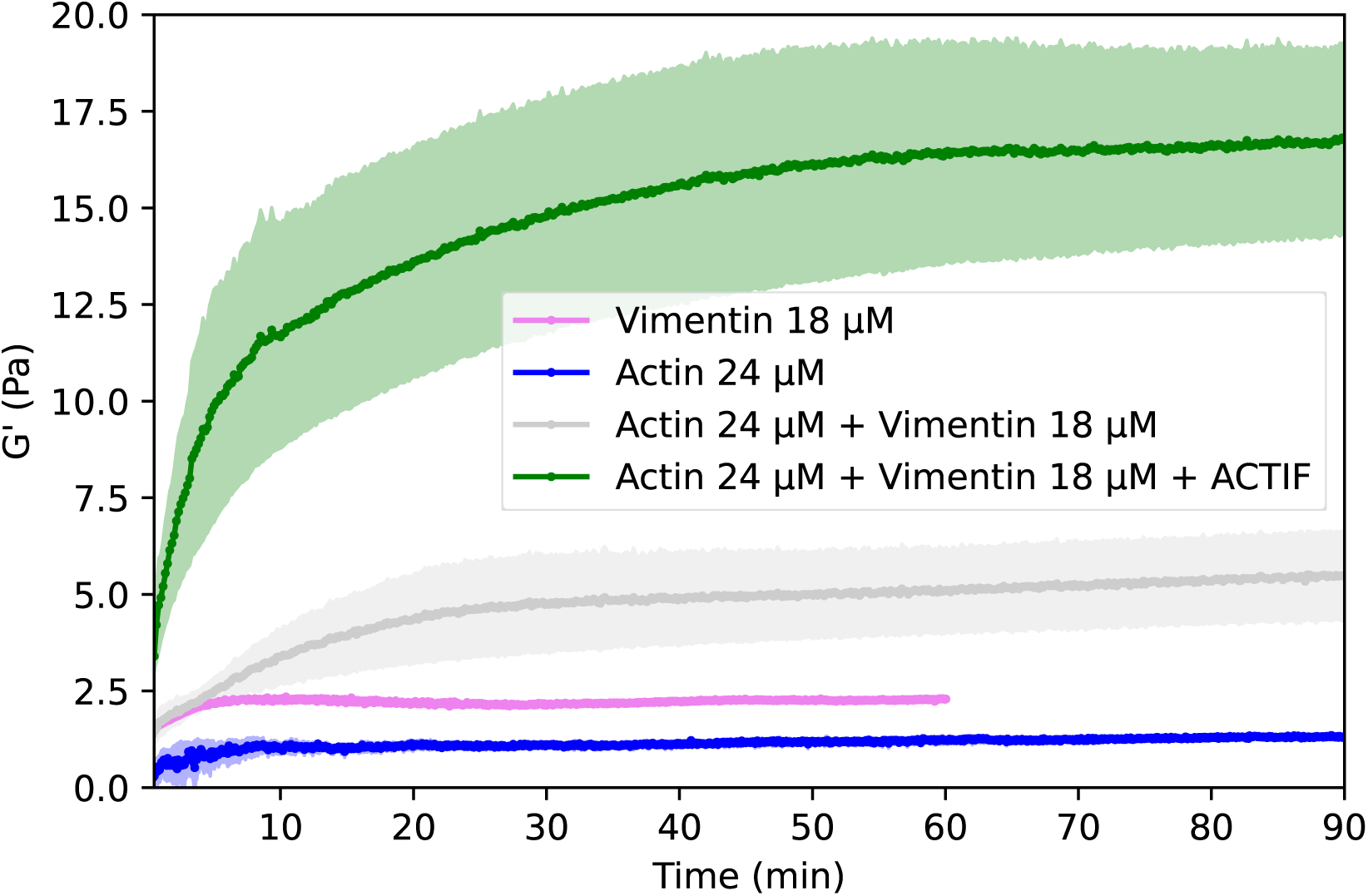
Polymerization curves for different reconstituted networks recorded by small amplitude oscillatory shear measurements. Actin (24 µM, blue), vimentin (18 µM, pink), actin and vimentin composite (24 µM actin, 18 µM vimentin, grey), and actin and vimentin composite crosslinked with 4.2 µM ACTIF (green). Curves averaged over N=2 measurements.

**Supplemental Figure 15:**
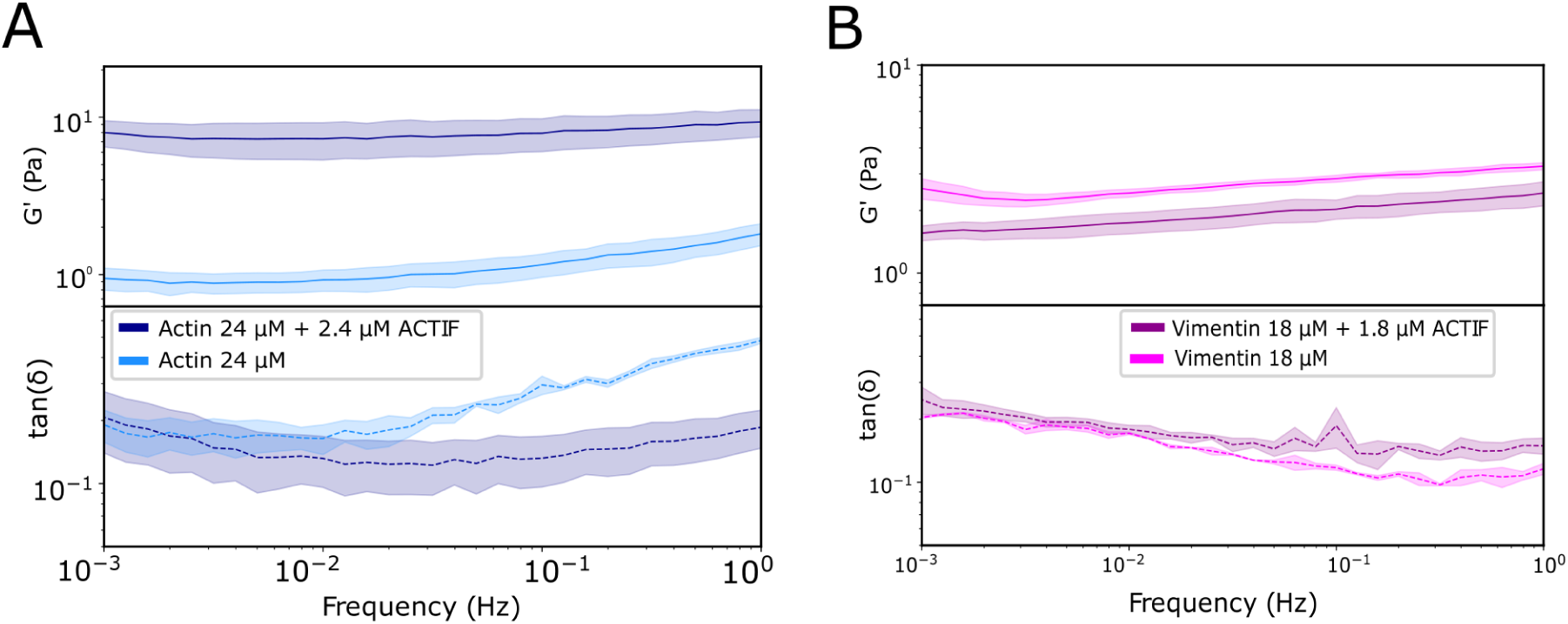
(A) Frequency sweep for an actin network (24 µM) and for a crosslinked actin network (24 µM actin + 2.4 µM ACTIF), showing the elastic and viscous shear moduli (top) and corresponding loss tangents (bottom). (B) Frequency sweep for a vimentin network (18 µM) and for a crosslinked vimentin network (18 µM + 1.8 µM ACTIF) showing the elastic and viscous shear moduli(top) and corresponding loss tangents (bottom).

**Supplemental Figure 16:**
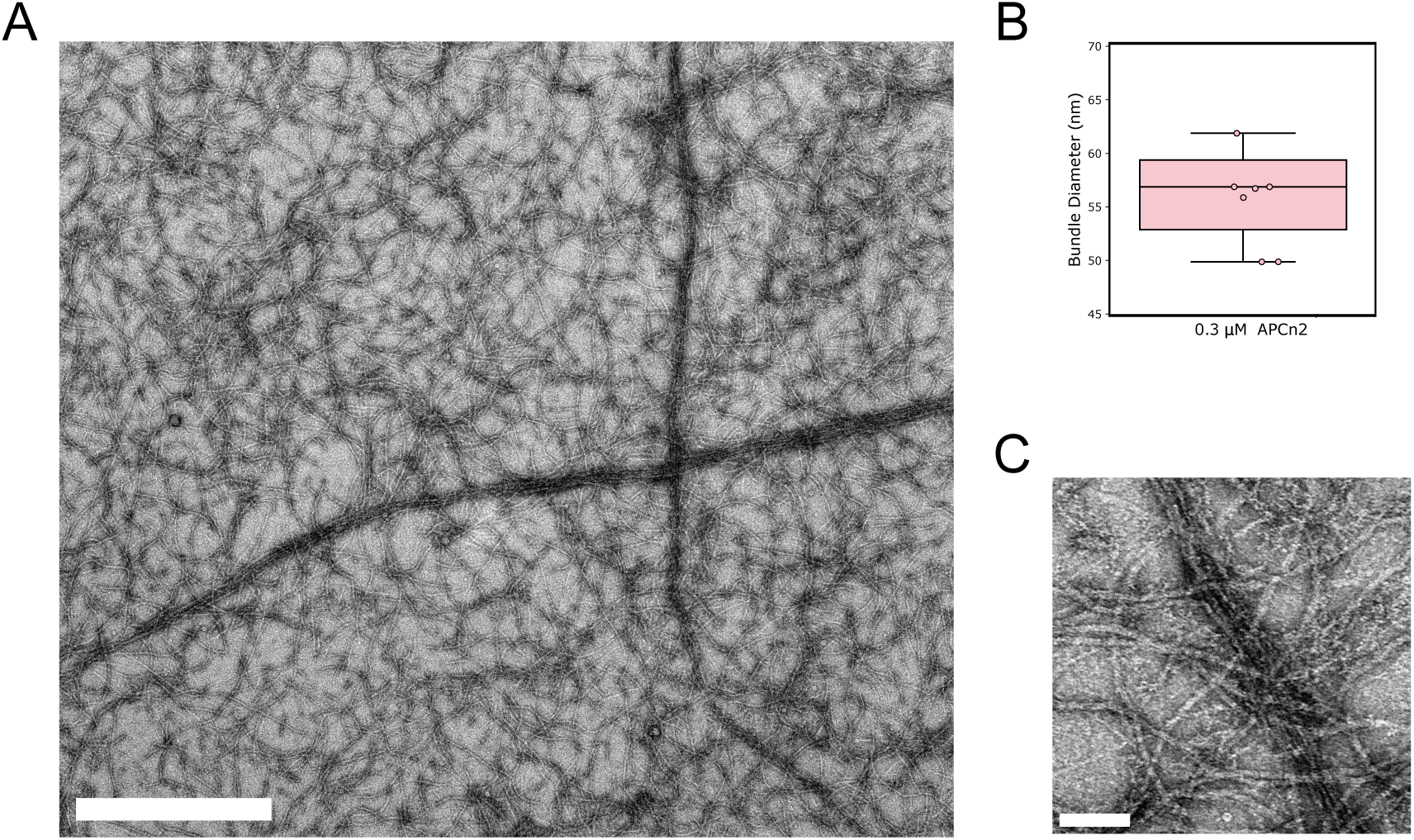
Transmission electron microscopy images of bundles of F-actin and vimentin formed with ACTIF-APCn2. (A) Overview of copolymerized actin (1.5 µM) and vimentin (1.5 µM) bundles formed with 0.3 µM of ACTIF-APCn2. Scale bar 1 µm. (B) Box plot depicting the bundle widths of 7 bundles from 2 independently prepared EM grids. (C) Zoomed-in image of a bundle. Scale bar 100 nm.

**Supplemental Figure 17:**
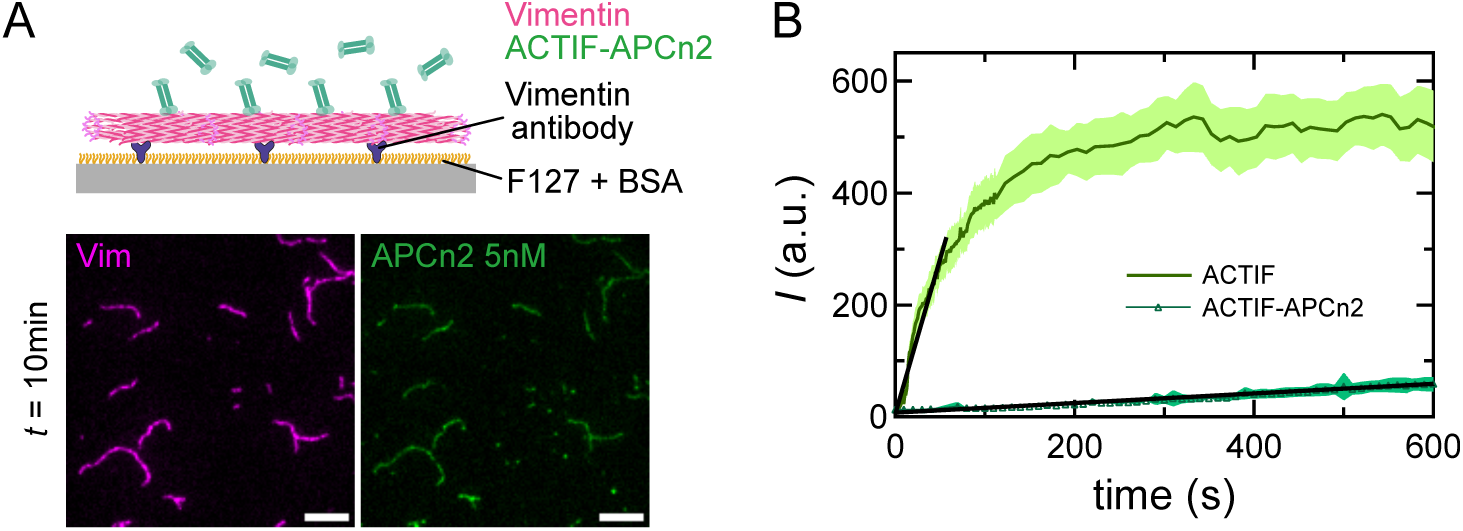
Binding affinity of ACTIF-APCn2 for vimentin filaments. (A) Schematic of the binding assay and images of ACTIF-APCn2 (green) at 5 nM co-localized with vimentin filaments (magenta) attached to the substrate, at *t* = 10 min after flushing ACTIF-APCn2 into the chamber. (B) Fluorescence intensity of ACTIF-APCn2 at 5 nM (dark green), in comparison with ACTIF at the same concentration of 5 nM (light green), on vimentin filaments over time (averaged over 30 filaments). Black lines represent linear regression of the curves. The shaded area represents standard deviations. Scale bar: 5 µm.

**Supplemental Figure 18:**
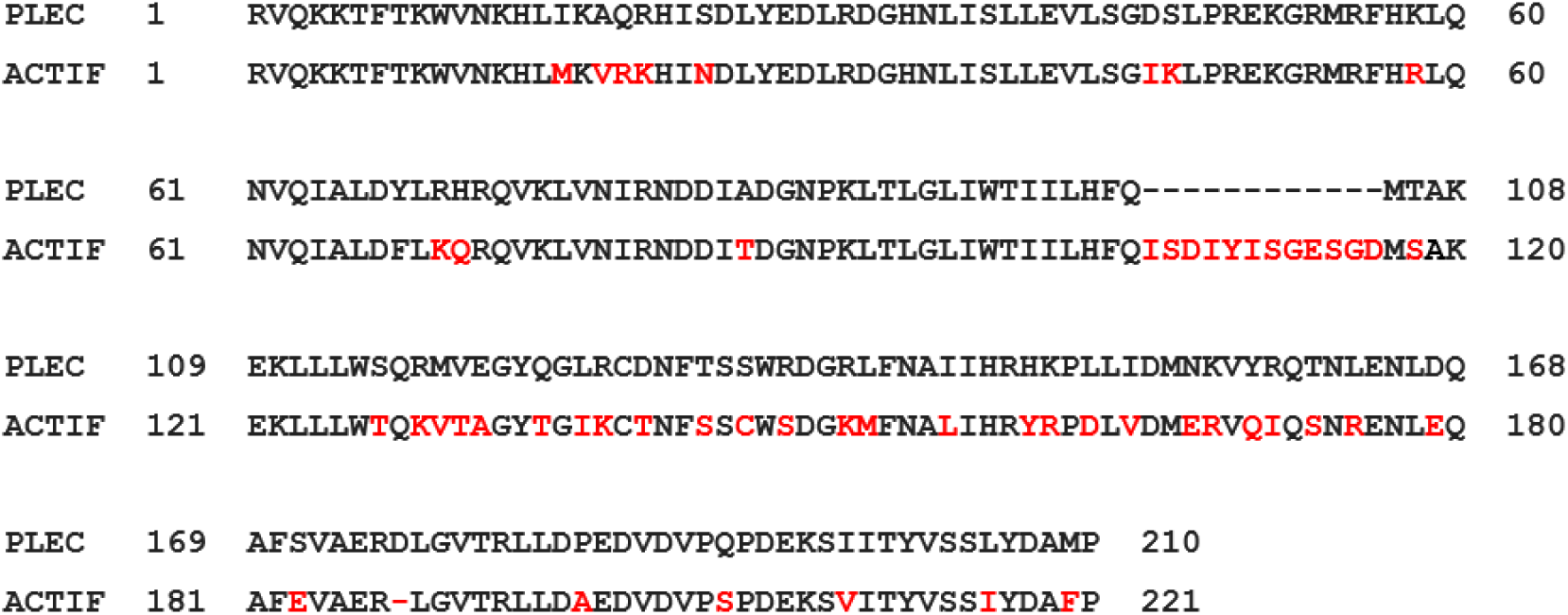
Sequence alignment of the calponin homology domains of plectin (top) and ACTIF (bottom), where the CH tandem of ACTIF has been taken from MACF (Q9QXZ0, 78-298 aa). In red, sequence differences.

**Supplemental Figure 19:**
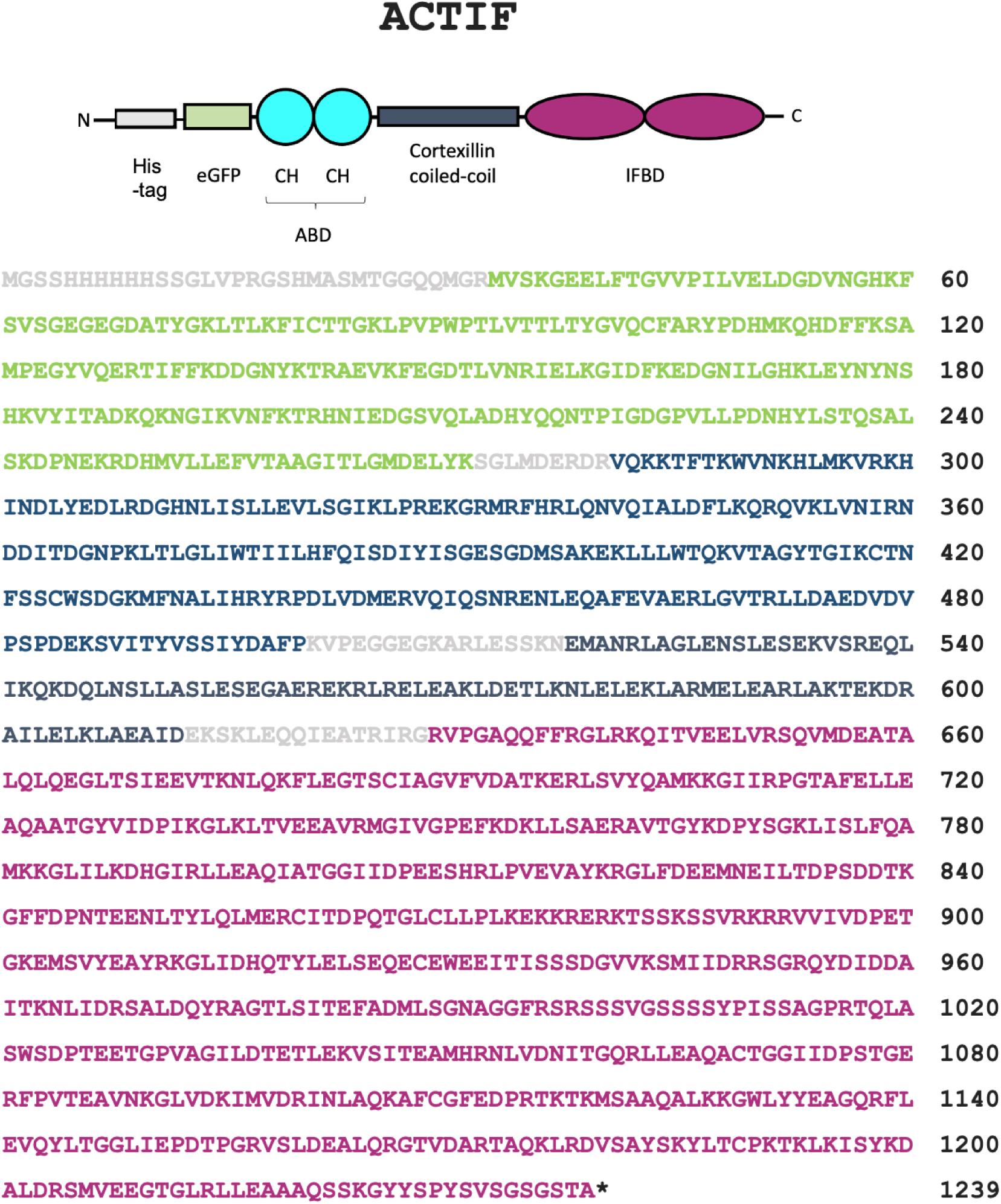
ACTIF’s amino-acid sequence and domain organization. (A) ACTIF domain organization with the amino-acid numbers indicated from the sequence in (B). eGFP, enhanced green fluorescent protein; ABD, actin-binding domain; CC, coiled-coil; IFBD, intermediate filament binding domain (B) Amino-acid sequence of ACTIF. The color code corresponds to the domains indicated on the diagram in (A) and the spacers between domains are indicated in light grey letters.

**Supplemental Figure 20:**
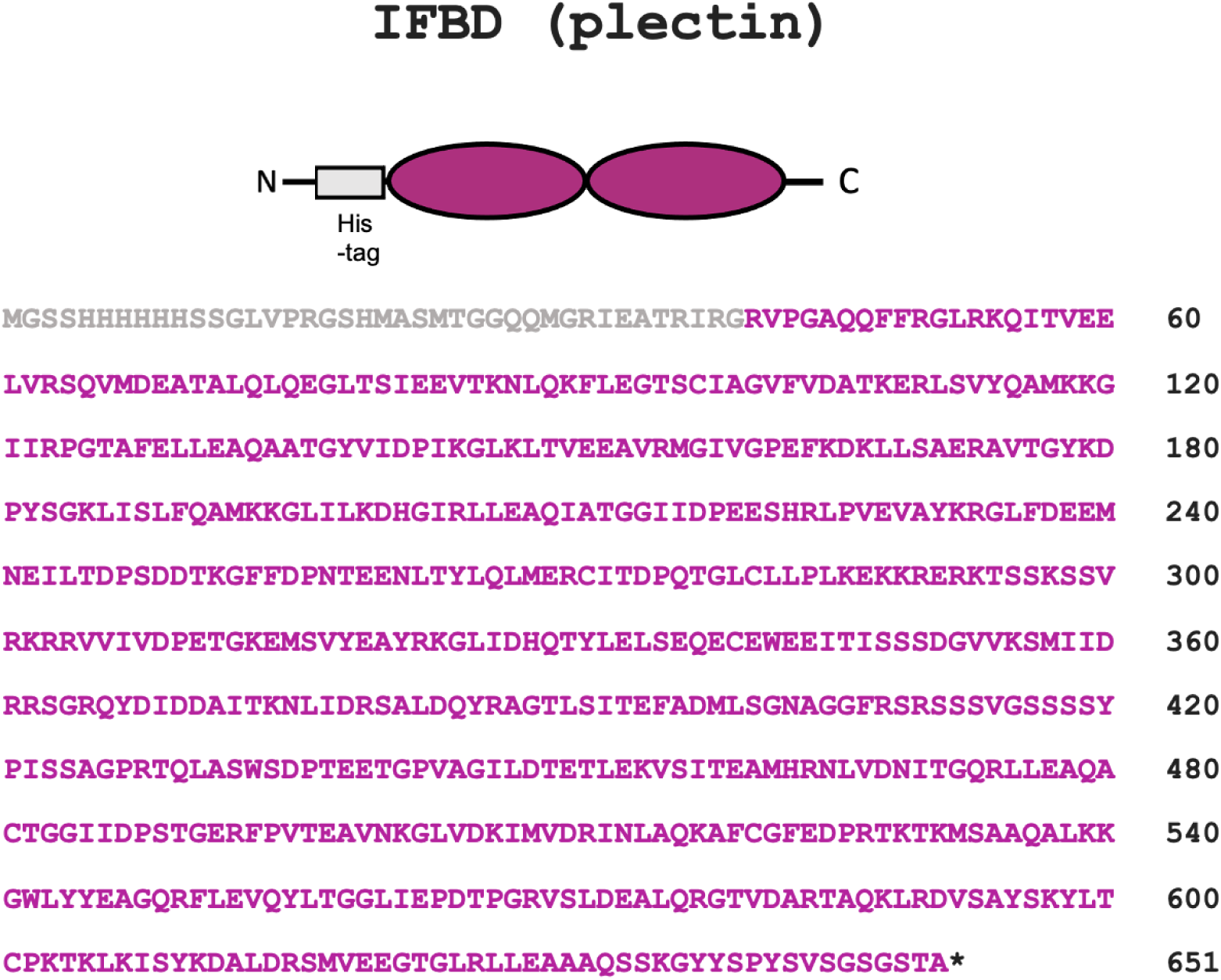
IFBD’s domain organization and amino-acid sequence. (A) Domain organization of IFBD (plectin). (B) Amino-acid sequence of IFBD (plectin) domain. The color code corresponds to the domains indicated on the diagram in (A).

**Supplemental Figure 21:**
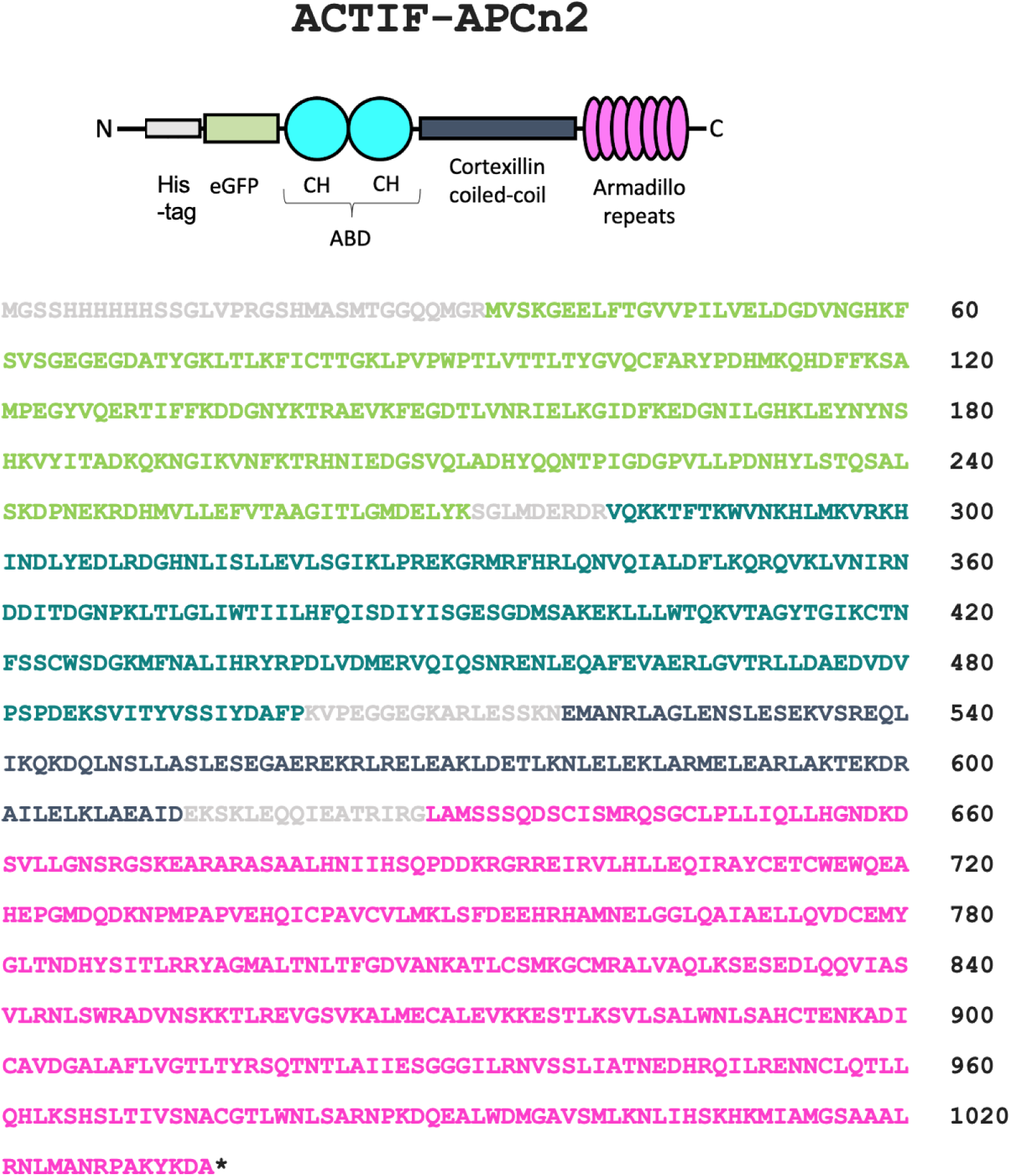
ACTIF-APCn2’s amino-acid sequence and domain organization. (A) Amino-acid sequence of ACTIF-APCn2. The color code corresponds to the domains indicated on the diagram in (B) and the spacers between domains are indicated in light grey letters. (B) ACTIF-APCn2 domain organization with the amino-acid numbers indicated from the sequence in (A). His-tag, grey; eGFP, enhanced green fluorescent protein; ABD, actin-binding domain; CC, coiled-coil; armadillo repeats.

### SUPPLEMENTARY MOVIE CAPTIONS

**Supporting movie 1**

Control experiment by TIRF microscopy showing no direct interaction between vimentin and actin filaments in our experimental conditions. Pre-assembled vimentin filaments (magenta) were stabilized on the coverslip substrate coated with vimentin antibodies, while F-actin (cyan) was growing from G-actin at 1 µM in F-buffer supplemented with 0.2 % methyl-cellulose, and moving freely on top of vimentin filaments with no signs of co-localization for 20 min.

**Supporting movie 2**

TIRF microscopy experiment to probe ACTIF-mediated binding of F-actin to vimentin filaments. First, ACTIF (green) at 5 nM in F-buffer was flushed into the chamber containing settled pre-assembled vimentin filaments (magenta) on the substrate for 10 min. Then, pre-assembled F-actin at 0.1 µM in F-buffer, without the addition of ACTIF, was flushed into the chamber. F-actin rapidly attached and co-localized with ACTIF-bound vimentin filaments.

